# Characterizing mitochondrial copy number variation and PCR amplification bias as sources of quantitative constraints in DNA metabarcoding

**DOI:** 10.64898/2026.04.07.716896

**Authors:** Lisa Wolany, Kevin Klinkenborg, Florian Leese, Dominik Buchner

## Abstract

DNA metabarcoding is a central tool in biodiversity research and monitoring, producing detailed taxa lists with comparatively little time and effort. One of its limitations, however, is the lack of quantitative data on biomass or abundance. This limitation has two main reasons: 1) DNA template copy number variation and 2) primer-induced PCR amplification bias. Many metabarcoding markers are mitochondrial and mitochondrial DNA copy numbers vary in animal tissues, potentially decoupling sequence counts from biomass. Additionally, primer mismatches can lead to taxon-specific amplification biases, for which PCR cycle calibration has been proposed as a solution. To mechanistically study both effects, we constructed and analyzed mock communities of different arthropod species. We combined digital PCR and COI metabarcoding to quantify relationships between biomass, mitochondrial copy number, and metabarcoding reads. Mitochondrial DNA copy numbers per biomass varied strongly within and among the different taxa. Metabarcoding reads did not reflect input mitochondrial DNA copies without a correction. Attempts to correct for amplification bias via PCR cycle calibration failed as read proportions remained stable across cycles. We therefore mathematically derived an approach to estimate relative amplification bias and initial mitochondrial DNA copy numbers in a sample based on a non-exponential amplification bias model and demonstrate its applicability. Still, the detected high variation in mitochondrial copy numbers and derived prerequisites necessary to calculate amplification efficiencies and mitochondrial copy numbers limit the practical application. Our study highlights fundamental constraints of quantitative metabarcoding and underscores the need for additional methodological approaches for quantitative insights while delivering essential conceptual insights.

## Introduction

DNA metabarcoding has revolutionized biodiversity research and monitoring (Abrego et al., 2024; Taberlet et al., 2012). The method delivers highly resolved taxa lists even for species-rich samples at moderate costs (Buchner et al., 2025). Consequently, several large-scale monitoring programs already employ DNA metabarcoding (Gilbert et al., 2014; Hardwick et al., 2024; Miraldo et al., 2025). Despite these advantages, a major challenge of the method remains the reliable estimation of biomass or abundance from sequence read data (Elbrecht & Leese, 2015; Sickel et al., 2023). Quantitative data, however, are essential for numerous ecological applications including many biodiversity indices and ecological status class assessment in biomonitoring frameworks such as the European Water Framework Directive (Hering et al., 2018). Like all high-throughput sequencing data, DNA metabarcoding outputs are compositional, i.e. the resulting read counts reflect relative abundances at best and cannot be directly related to the number of DNA template molecules in the original sample (Gloor et al., 2017). Therefore, metabarcoding sequence reads cannot be used as a direct replacement for traditional abundance or biomass information. Furthermore, the interpretation of inferred relative read abundances is complicated by several factors that distort the relationship between input DNA template molecules and observed reads.

A first potential source of bias is the number of copies of the target marker gene in the sample, which for many animal target groups is the copy number of mitochondrial DNA marker genes (mtDNA copies), such as cytochrome c oxidase subunit I (COI), cytochrome b, 16S rDNA, and 12S rDNA (deWaard et al., 2019; Hebert et al., 2003; Karlsson et al., 2020). Consequently, the number of amplifiable marker target molecules in a sample depends not only on organism biomass but also on the number of mtDNA copies present in the tissue. While the variation in mtDNA copy number is widely recognized as a potential source of bias (Krehenwinkel et al., 2017; Piñol et al., 2019; Shaffer et al., 2025), it has received relatively little attention in the DNA metabarcoding community. In contrast, studies on human and murine tissue have demonstrated that mtDNA copy number can vary as much as 200-fold among and within tissues (Rath et al., 2024). Similarly, chloroplast DNA copy number – the analogous organellar target in plant metabarcoding – has been shown to vary up to 67-fold between tissue types even within the same species (Kamenova et al., 2025). This suggests that copy-number variation in organellar marker genes is a widespread, challenge across taxa for quantitative metabarcoding. A prerequisite for reliably inferring quantitative information from sequence reads in complex samples is therefore either the correction for species-specific variation in mtDNA copy number, or the assumption that interspecific variation in mtDNA copy number is negligible. Crucially, however, any such correction is only meaningful if mtDNA copy numbers scale predictably with organismal biomass. If this relationship is weak or highly variable, the reliability of any quantitative inference from metabarcoding reads will be substantially compromised.

In addition to mtDNA copy number variation, PCR amplification bias, i.e. the preferential amplification of certain DNA template molecules during PCR, can further distort quantitative relationships. The kinetics of PCR amplification are commonly described by an exponential relationship:

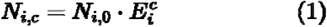

where ***N***_***i*,*c***_ represents the number of amplified DNA sequences for taxon ***i*** after ***c*** cycles, ***N***_***i*,0**_ the initial number of target DNA templates for taxon ***i*** and ***E***_***i***_ represents the taxon-specific amplification efficiency, where ***E* = 1** denotes no net amplification and ***E* = 2** perfect doubling per cycle (Ruijter et al., 2009; Shelton et al., 2023; Silverman et al., 2021).

Mismatches between primer and target sequence have been identified as a key driver of variation in this taxon-specific amplification efficiency (Shaffer et al., 2025; Shelton et al., 2023). Silverman et al. (Silverman et al., 2021), however, argue that primer– template mismatch effects are confined to the first three cycles of PCR, after which the original template at the primer binding site is replaced by the sequence of the introduced primer. Therefore, they attribute the substantial bias observed in mid-to-late cycles to non-primer-mismatch sources, such as choice of Taq polymerase, sequence length, and GC content (Nichols et al., 2018). Notably, both explanations employ the same exponential amplification model and agree on the empirical consequence: small differences in ***E***_***i***_ between taxa, irrespective of their mechanistic source, are compounded across cycles and translate into large systematic biases in final read proportions.

While the preceding considerations apply to any single primer, degenerate primers introduce an additional layer of complexity as they consist of a mixture of several primers. In such mixtures, ***E***_***i***_ is not the efficiency of a single primer–template interaction but rather the composite binding efficiency of the subset of primer variants that successfully anneals to each taxon’s target DNA fragment — an emergent property of the mixture that differs between taxa and cannot be reduced to any single molecular interaction. However, as many studies target phylogenetically diverse biological samples, such as those collected from soil, aquatic environments, or even terrestrial insect traps, the diversity of target DNA templates is extremely high, making the use of such degenerate ‘universal’ primers inevitable. Especially for protein-coding marker genes, like COI, primers are often degenerate due to codon redundancy. As an example, for 23-26 bp primers like the fwh2/fwhR2n primer pair (Vamos et al., 2017), this comprises more than 200 individual primer sequences in one degenerate primer mixture. For any given taxon, the subset of these variants that successfully anneals — and their combined binding efficiency — constitutes the effective ***E***_***i***_, which is thus a difficult-to-predict quantity (Piñol et al., 2019).

Furthermore, neither ***N***_***i*,*c***_ nor ***E***_***i***_ can be directly observed or estimated due to the compositional nature of sequencing data. Instead, what is empirically accessible from a sequencing run is the relative read abundance ***R***_***r***_ of each taxon, i.e. its share of the total sequenced amplicon pool at a given cycle ***c***:

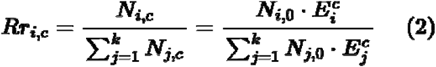

This share is not stable across cycles, since ***E***_***i***_ typically differs among taxa. During the exponential phase of PCR, it shifts progressively: Taxa with higher efficiencies become overrepresented, while taxa with lower efficiencies become underrepresented. Thus, it becomes apparent, that the relative read abundance of any taxon is influenced by the amplification efficiencies of all other taxa in a sample.

Various strategies have been proposed to mitigate amplification bias in metabarcoding data. One involves deriving species-specific correction factors from artificially constructed communities (‘Mock community strategy’; (Darby et al., 2020; Shelton et al., 2023; Thomas et al., 2016). A limitation of this strategy is the requirement to derive a correction factor for every species contributing to the observed community. This, however, presents a considerable challenge for highly diverse groups such as insects, as typically a non-negligible portion of the samples’ diversity is either unknown (no correction factors can be inferred) or not amplified.

An alternative strategy, which allows for the adjustment of community-specific bias, estimates the amplification efficiency for each specific taxon and community by varying the number of PCR cycles (‘Cycle calibration strategy’, (Shelton et al., 2023). The PCR cycle calibration strategy utilizes measurements of the produced DNA sequences following different numbers of PCR cycles to calculate the initial starting concentration of their respective DNA templates (Kelly et al., 2019; Silverman et al., 2021). An apparent advantage of this correction method is that ***E***_***i***_ values and therefore starting concentrations can be inferred for any given sequence in a sample. However, empirical support for this bias correction strategy is mixed. While Silverman et al. (Silverman et al., 2021) successfully demonstrated that the method mitigated PCR bias, Shelton et al. (Shelton et al., 2023) reported a lack of success employing the method, primarily attributing this to the challenge of generating sufficient sequenceable amplicon with a restricted number of PCR cycles. Also, Krehenwinkel et al. (Krehenwinkel et al., 2017) indicated that reducing the cycle number did not influence amplification bias. Consequently, the reliability and general applicability of PCR cycle calibration across all sample types remains uncertain.

In this study we aimed to investigate key prerequisites for generating quantitative data from metabarcoding reads when using mitochondrial marker genes. First, we examined the relationship between biomass and mtDNA copy number and subsequently tested if through PCR cycle calibration individual amplification efficiencies can be derived. To achieve this, we constructed and analyzed a set of 81 mock communities with systematic ranges of biomass and species composition from five arthropod species. MtDNA copy numbers of all samples were quantified using digital PCR (dPCR) and DNA metabarcoding was performed for all mock communities using the cycle calibration strategy. To better understand the underlying molecular mechanics, we replicated DNA metabarcoding analyses with a second primer set targeting the same marker. Additionally, we used our findings to derive a conceptual mechanistic framework for PCR amplification bias that may inform future correction strategies.

We hypothesized that:

1. Species biomass proportions are reflected in relative mtDNA copies.
2. Uncorrected relative read numbers from metabarcoding analyses do not reflect the relative copy number of template sequences.
3. PCR cycle calibration enables estimation of taxon-specific amplification efficiencies within the studied community of interest.
4. Correcting relative read numbers for amplification bias enables accurate estimation of relative template copy numbers.

## Materials and Methods

### Workflow overview

We constructed mock communities from five arthropod species of different species compositions and biomass ratios. The samples were homogenized and two aliquots were taken from each sample (Figure 1). DNA from each aliquot was extracted in duplicates. After DNA extraction, mtDNA copy numbers of replicates A and B were measured via digital digital PCR (dPCR). Metabarcoding was performed for all replicates of both aliquots and only sequences present in both replicates of one aliquot were considered valid. Taxon-specific amplification biases were quantified using calibration communities of known composition and used to convert metabarcoding read proportions into estimates of relative mtDNA copy numbers.

**Figure 1.**
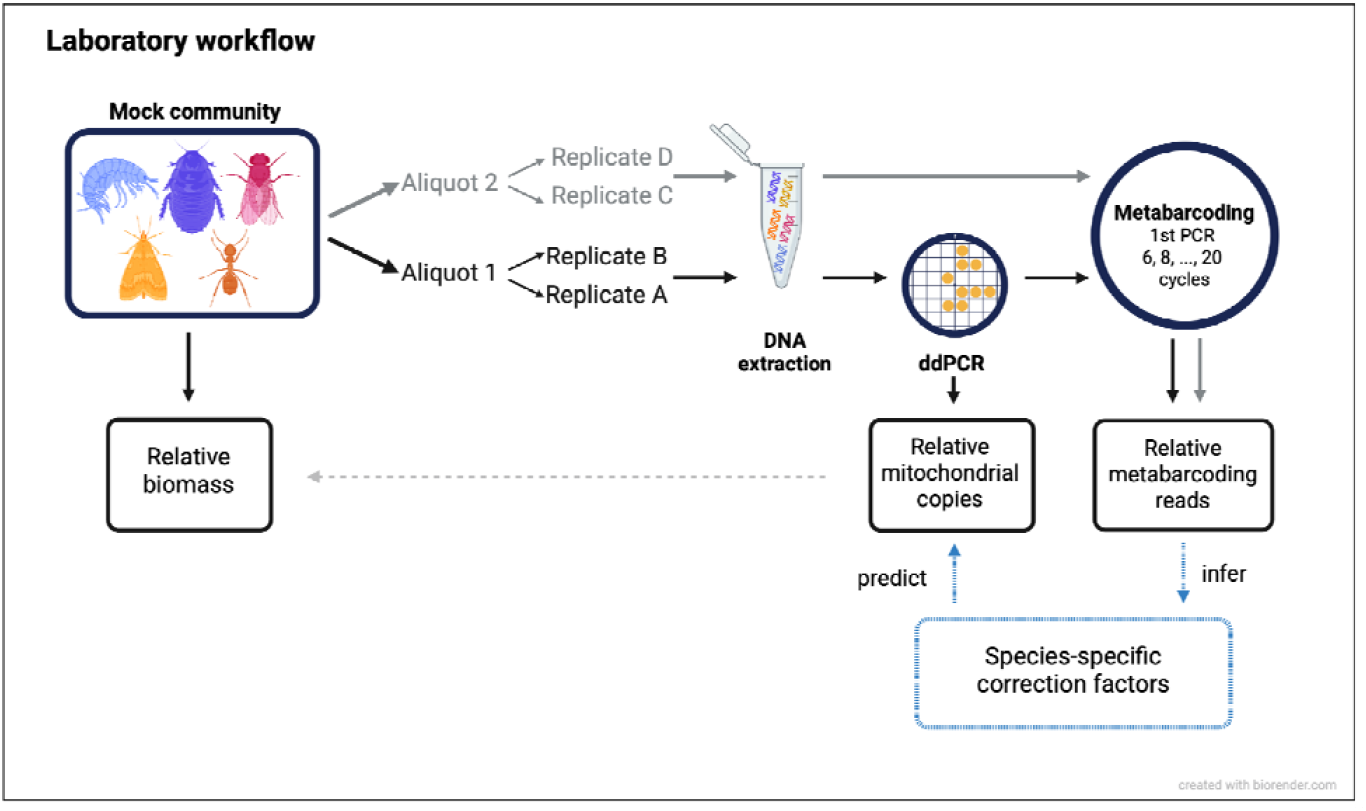
Overview of study workflow.

### Mock community assembly and DNA extraction

Mock communities were constructed from five arthropod taxa that will hereafter be referred to at order level, i.e.: Hymenoptera: *Acromyrmex* sp. (Mayr, 1865), Blattodea: *Blaptica dubia* (Serville, 1838), Lepidoptera: *Galleria mellonella* (Linnaeus 1758), Amphipoda: *Gammarus* sp. (Fabricius, 1775), and Diptera: *Hermetia illucens* (Linnaeus 1758). The animals were obtained as live specimens from live feed supply and hobbyist animal cultures. All individuals belonging to one species came from one supply to guarantee their genetic similarity. The animals were preserved in 96% methylated ethanol, with an ethanol change 28-48 hours after initial preservation.

All individuals were dried in a vacuum centrifuge and weighed subsequently. Mock communities were constructed so that all possible combinations of two-, three-, four-, and five species communities were considered. For each community type, a gradient of biomass ratios ranging from 5% to 90% percent of sample biomass per species was covered in up to 5 replicates (see supporting information 1). 84 communities were assembled to achieve a full PCR-plate setup in combination with 12 negative control samples. Three samples were excluded from the dataset after sequencing revealed that they had been incorrectly assembled during sample preparation, resulting in a community composition that did not match the intended design. This left 81 samples for the final analysis.

After assembling the communities from individual specimens, the samples were filled up to a volume of 30 mL with 96% ethanol and subsequently homogenized with a rotor-stator-homogenizer (Tissue-Tearor, Biospec Products Inc., Bartlesville, OK, USA). Before DNA extraction 12 negative controls of pure ethanol (96%) were added to the pool of samples, so that every row and column in a 96-well plate contained at least one negative control. All DNA extraction and metabarcoding steps were performed on an automated liquid handler (Hamilton Microlab Vantage, Hamilton Bonaduz AG, Bonaduz, Switzerland), according to the workflow recommendations from Buchner et al. (Buchner et al., 2021) and following our standard protocol available on protocols.io (Buchner, 2022a).

1 mL of homogenized tissue was used for sample lysis. Sample lysis was performed by removing the ethanol and subsequent addition of 900 µL lysis buffer and 100 µL Proteinase K (10 mg/mL) (Buchner, 2022d). Bead-beating was followed by 20-minute incubation at 56 °C at 1,400 rpm. Extraction replicates were structured as follows: A 1 mL aliquot of every sample was used to create extraction replicates A and B; while extraction replicates C and D were taken from another 1 mL aliquot of the sample homogenate. Subsequently, DNA was extracted with a silica-bead-based protocol (Buchner, 2025a). The success of the DNA extraction was controlled on a 1%-agarose gel.

### dPCR probe design

Species-specific primers and probes were designed for each of the five target species with Primer3 (Untergasser et al., 2012), with probes labeled with a 5’ FAM reporter and a 3’ TAMRA quencher and tested in silico to avoid co-amplification among target species (Supporting information 2). Primers and probes were synthesized by Eurofins Genomics (Eurofins Genomics Europe Shared Services GmbH, Ebersberg, Germany). Assay specificity was subsequently validated by qPCR, confirming correct amplification of target species and the absence of amplification for non-target species within each community (Supporting information 3, Supporting information 4).

### dPCR measurements

dPCR was performed on a QuantStudio Absolute Q Digital PCR System. Reagents used per reaction were 2 µL Absolute Q Universal DNA Digital PCR Master Mix (5X) (Thermo Fisher Scientific, Darmstadt, Germany) and 6 µL of assay premix containing 1000 nM of each primer and 750 nM probe, with the following protocol: 10 min at 96 °C followed by 40 cycles of 98 °C for 15 s and 60 °C for 30 s. DNA extracts were measured at a 1:1000 dilution. For samples with insufficient precision reported by the instrument, measurements were repeated at a 1:100 dilution to achieve precision levels within 10% (Supporting information 5).

### Metabarcoding

Metabarcoding was performed for all 4 extraction replicates (plates A-D) using a two-step PCR protocol (Zizka et al., 2019). For the investigation of the cycle calibration strategy, 8 different PCR cycling thresholds were applied to the replicates of each plate. Cycle numbers of the first-step PCR were chosen in 2-cycle increments between 6-20 cycles, resulting in one PCR replicate each with 6, 8, 10, 12, 14, 16, 18 and 20 1st-step PCR cycles per initial extraction replicate plate.

For the 1st step PCRs with different cycle numbers the fwhF2/fwhR2n primer pair (fwhF2: 5’-GGDACWGGWTGAACWGTWTAYCCHCC-3’, fwhR2n: 5’-GTRATWGCHCCDGCTARWACWGG-3’, (Vamos et al., 2017), hereafter “fwh2”, was used, with the following PCR conditions: 10 min of initial denaturation at 95 °C, 30 s denaturation at 95 °C, 90 s annealing at 58 °C, 30 s elongation at 72 °C and a final elongation of 10 min at 68 °C, whereas the steps denaturation-elongation were repeated for the respective cycle number. Subsequently, a size selection of the PCR products was carried out with carboxylated beads (Buchner, 2022b). The second PCR step was conducted with primers that bind to a universal binding site attached through the first PCR primer (forward universal tail: 5’-ACACTCTTTCCCTACACGACGCTCTTCCGATCT-3’, reverse universal tail: 5’GTGACTGGAGTTCAGACGTGTGCTCTTCCGATCT-3’, (Werner & Buchner, 2025), leading to an equal amplification efficiency for all sequences during the second PCR step. The cycle number for the second PCR step was chosen in relation to the cycle number of the first PCR for every replicate, to achieve a sum of 45 PCR cycles in total, congruent with our standard established metabarcoding workflow (Buchner et al., 2025). After the 2nd-step PCR, successful amplification PCR was verified on a 1% agarose gel. The individually tagged samples were subsequently normalized and size selected with carboxylated beads (Buchner, 2022c) and pooled into two libraries. The libraries were concentrated with silica spin columns and the DNA concentration measured with a Qubit fluorometer (Thermofischer, Darmstadt, Germany). A reconditioning PCR was performed for both libraries to reduce PCR-associated artifacts such as heteroduplexes and chimeras. This involved a short, one-cycle reamplification with fresh reagents, using the flow cell binding adapters P5 and P7 as primers (Buchner, 2025b). Replicates of all 4 plates were also amplified with the BF3/BR2 primer set (BF3: 5’-CCHGAYATRGCHTTYCCHCG-3’, BR2: 5’-TCDGGRTGNCCRAARAAYCA-3’, Elbrecht et al., 2019), hereafter “BF3”, with 20 cycles of first PCR and 25 cycles of second PCR at identical conditions as stated above except for an annealing temperature of 50 °C in first PCR and sequenced in one library. Sequencing was performed on an Illumina NovaSeq platform by Genewiz/Azenta (Leipzig, Germany) for the fwh2 libraries and by CeGaT (Tübingen, Germany) for the BF3 library.

### Bioinformatic processing

The metabarcoding raw data were demultiplexed via the python package demultiplexer2 (v 1.1.4, https://github.com/DominikBuchner/demultiplexer2). Subsequent data processing (paired-end merging, primer trimming, quality filtering, dereplication, denoising, replicate merging, removal of negative controls) into a read table was performed via the APSCALE pipeline (v. 4.3.0, (Buchner et al., 2022), default settings). The taxonomic assignment of the resulting sequences was conducted with BOLDigger3 (v. 3.0.2, (Buchner & Leese, 2020). The raw pipeline output for both primers, including taxonomic assignments and sample metadata, is provided in supporting information 6. After taxonomic assignment reads were pooled at order level. Data were filtered to retain only the taxa expected in each sample, which removed 0.07% of reads in the fwh2 dataset and 0.78% of reads in the BF3 dataset. To test for homogeneity of metabarcoding aliquots and dPCR replicates, corresponding sets were plotted against each other (supporting information 7, supporting information 8) and no significant differences between replicates were detected using a Wilcoxon signed-rank test. The measurements of mtDNA copy number from the dPCR were averaged for replicates A and B. After verifying replicate consistency, metabarcoding replicates were pooled. In the metabarcoding dataset, an entry of zero reads was created for all taxa that were part of the sample but not detected by metabarcoding. As all measurements were conducted on multi-species samples, proportions of mtDNA copy numbers and metabarcoding reads were calculated by dividing the individual value for every species by the sum of mtDNA copies or reads of the sample. Biomass data were expressed as taxa proportions of the total sample biomass.

To quantify how accurately biomass proportions were reflected by mtDNA copy numbers and metabarcoding reads, we computed log_2_ fold-changes between all three pairwise comparisons: relative biomass vs. relative mtDNA copies, relative mtDNA copies vs. relative metabarcoding reads, and relative biomass vs. relative metabarcoding reads. For each species in a sample, the fold-change was calculated as:

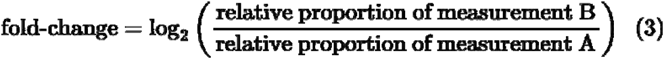

A fold-change of 0 indicates perfect agreement between two measurements, while positive values indicate overrepresentation and negative values indicate underrepresentation in measurement B relative to A. The log_2_ scale ensures symmetry: a value of +1 corresponds to a 2-fold overrepresentation, −1 to a 2-fold underrepresentation. Species with zero reads were excluded from the calculation. To assess agreement, we defined a 1.5-fold tolerance band (log_2_ ≈ ±0.585) and reported the proportion of species falling within this band for each comparison. Additionally, we assessed rank concordance by computing the proportion of species that retained their within-sample rank across measurement types.

PCR cycle calibration was performed as suggested by Shelton et al. (Shelton et al., 2023) by fitting, for each sample, a linear regression of the log read-count ratio between each taxon and an arbitrarily chosen reference taxon against PCR cycle number. Predicted relative initial copy numbers were then reconstructed from the fitted intercepts via a softmax transformation.

To additionally evaluate amplification bias correction via mock communities as proposed by Shelton et al. (Shelton et al., 2023), we used the 14 two-species samples each containing Blattodea and one other taxon as calibration communities. For each calibration sample, the taxon-specific amplification efficiency per PCR cycle was estimated at ***c* = 20** as:

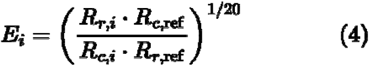

where ***R***_***r***_ and ***R***_***c***_ denote relative read and relative mtDNA copy proportions, respectively, and the subscript ref refers to Blattodea. Values were averaged across calibration samples. Predicted read proportions for the remaining 67 samples were obtained by dividing relative reads by 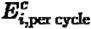, where ***c*** is the PCR cycle count of the respective sample.

Model performance was assessed using the coefficient of determination

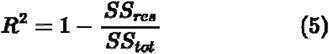

where ***SS***_***res***_ is the residual sum of squares between observed and predicted relative copy proportions and ***SS***_***tot***_ is the total sum of squares around the observed mean (Kvålseth, 1985). Unlike the classical ***R***^**2**^for linear regression with intercept, this formulation is not bounded below at zero and can take negative values when model predictions are worse than the observed mean.

We additionally calculated the Aitchison distance (Aitchison, 1982; Shelton et al., 2023) between predicted and observed relative taxon proportions for each sample, computed as the Euclidean distance between centered log-ratio (CLR)-transformed proportions. A full results table including fold-changes and ranks is provided in Supporting Information 9. The script to reproduce the entire analysis presented in this paper, including the generation of the figures and supporting information has been published on GitHub (https://github.com/DominikBuchner/quantitative_metabarcoding) and is provided as supporting information 10.

## Results

### Sequencing results

The sequencing of the fwh2 libraries yielded 1,271,494,502 total reads with an average sequencing depth of 413,898 reads per sample. After bioinformatic processing including taxonomic filtering, 1,088,663,970 reads were retained for data analysis. The sequencing of the BF3 library yielded 741,085,198 total reads with an average sequencing depth of 1,929,909 reads per sample. After bioinformatic processing including taxonomic filtering 473,220,495 reads were retained. All replicates tested for consistency (dPCR, metabarcoding with fwh2 and metabarcoding with BF3) showed high congruence between replicate pairs (Wilcoxon signed-rank test, p_dPCR_=0.860, p_fwh2_=0.382, p_BF3_=0.623).

### Variation of biomass, mtDNA copy number, and metabarcoding reads

In contrast to our expectation outlined in hypothesis 1, the species biomass proportions were not well reflected in relative mtDNA copies. In the analysis of biomass and mtDNA copy ranks within the samples, out of 210 possible species-sample (supporting information 1) pairs, only 112 (53.3 %) had identical ranks, while the remaining ranks differed. The rank concordance for the relationship of mtDNA copies and metabarcoding reads was high, with 75.7 % for the fwh2 primer and 74.8 % for the BF3 primer. Within-sample rank concordance was lowest for the relationship of biomass and metabarcoding reads (fwh2: 44.3 %, BF3: 43.8 %) (Table 1).

**Table 1.** Within-sample rank concordance of biomass, mtDNA copies, and metabarcoding reads for the primer pairs fwh2 and BF3 at cycle 20. Values indicate the number of species-sample pairs in which the rank of a species was concordant between both measurements (n/total, %). As copies were inferred via dPCR no difference for biomass – copy ranks exist between fwh2 and BF3.

| Primer | Biomass – Copies | Copies – Reads | Biomass – Reads |
| --- | --- | --- | --- |
| fwh2 | 112/210 (53.3 %) | 159/210 (75.7 %) | 93/210 (44.3 %) |
| BF3 |  | 157/210 (74.8 %) | 92/210 (43.8 %) |

Analysis of the fold-change calculations showed that the discordance of biomass representation by mtDNA copy number was not evenly distributed across the different taxa: For the selected 1.5-fold deviation threshold, Amphipoda was underrepresented in 78 % of all species-sample pairs, while between 67.5 % and 43.2 % of species-sample pairs were within this range for the remaining orders. Diptera were overrepresented in 56.8 % of the species-sample pairs by more than 1.5-fold. Fold-changes ranged from 2.63 to −3.91 across all taxa (Figure 2A, supporting information 9, supporting information 11).

**Figure 2.**
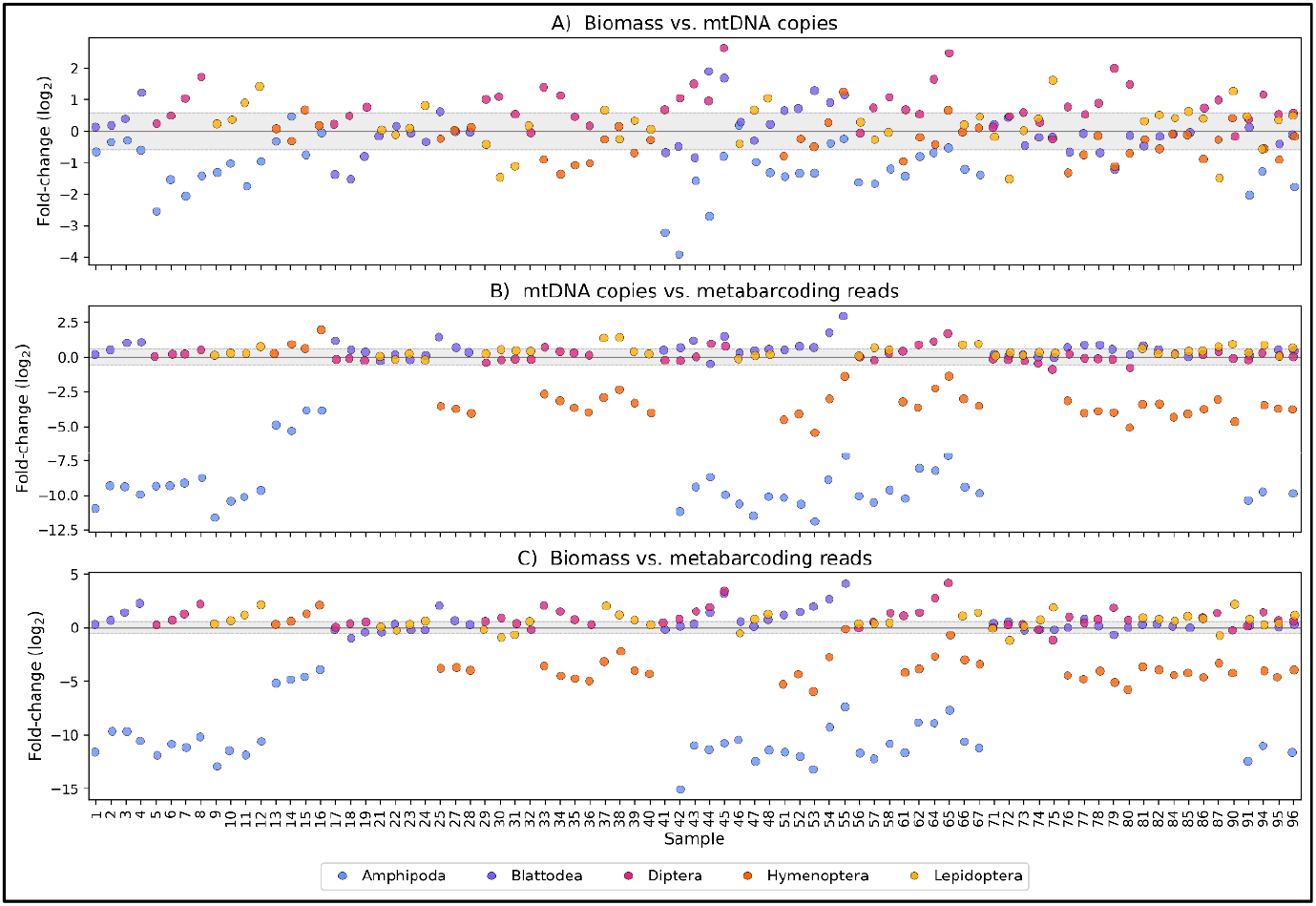
Per taxon fold-change between paired measurements of biomass, mtDNA copies and metabarcoding reads for all samples at cycle 20 (fwh2). Each point represents one species within one sample. The grey range indicates a 1.5-fold deviation threshold; points outside the range are classified as over- or underrepresented. **A)** Biomass vs. mtDNA copies. **B)** mtDNA copies vs. metabarcoding reads. **C)** Biomass vs. metabarcoding reads.

When comparing mtDNA copies and metabarcoding reads, the underrepresentation of Amphipoda became more pronounced, with Amphipoda being underrepresented in all samples. Hymenoptera were under-amplified as well, with 90.2 % of all species-sample pairs being underrepresented in metabarcoding reads compared to mtDNA copy numbers. Blattodea, Diptera, and Lepidoptera were least biased by the amplification. They were in concordance with the 1.5-fold change range in 63.6 %, 81.8 % and 72.5 % respectively. The comparison of mtDNA copy numbers and metabarcoding reads showed a greater variation than the comparison of biomass and mtDNA copies. Fold-changes ranged from 2.98 to −12.70 (Figure 2B, supporting information 9, supporting information 11).

The bias in representation became more pronounced when comparing biomass and metabarcoding reads, as the two biases are accumulated in this comparison. In the comparison of biomass and metabarcoding reads, the number of species-sample pairs within the 1.5-fold change threshold was the lowest (A: 107 of 210; B: 94 of 210; C: 63 of 210). Fold-changes ranged from 4.20 to −15.82 (Figure 2C, supporting information 9, supporting information 11). The effects were similar for the BF3 primer, with Hymenoptera showing the greatest bias (data not shown, supporting information 9). The per-sample community compositions for all samples and primers are shown in supporting information 12 and 13.

### PCR cycle calibration

Contrary to our expectation from hypothesis 3, the proportion of reads for each species and community did not show a systematic shift with an increasing number of PCR cycles (Supporting information 14). Consequently, the cycle calibration strategy, did not meaningfully improve the prediction of relative copy proportions compared to uncorrected reads (uncorrected vs. cycle calibrated, respectively: ***R***^**2**^ **= − 0.064** vs. ***R***^**2**^ **= − 0.039**; Figure 3, supporting information 15). Assuming that primer sequences are functionally replaced during the initial two cycles of amplification (as stated in (Silverman et al., 2021), all amplicons that have undergone a minimum of two PCR cycles, possess primer binding site sequences identical to the primer sequence that bound in the two preceding cycles (supporting information 16). This substitution effectively mitigates the systematic bias towards certain target sequences as a function of the number of PCR cycles. The principle applies to non-degenerate as well as to degenerate primers: As soon as the template sequence is functionally replaced by any of the primer variants, it perfectly binds to this specific variant, as long as there is excess primer, i.e. in the exponential phase of PCR.

**Figure 3.**
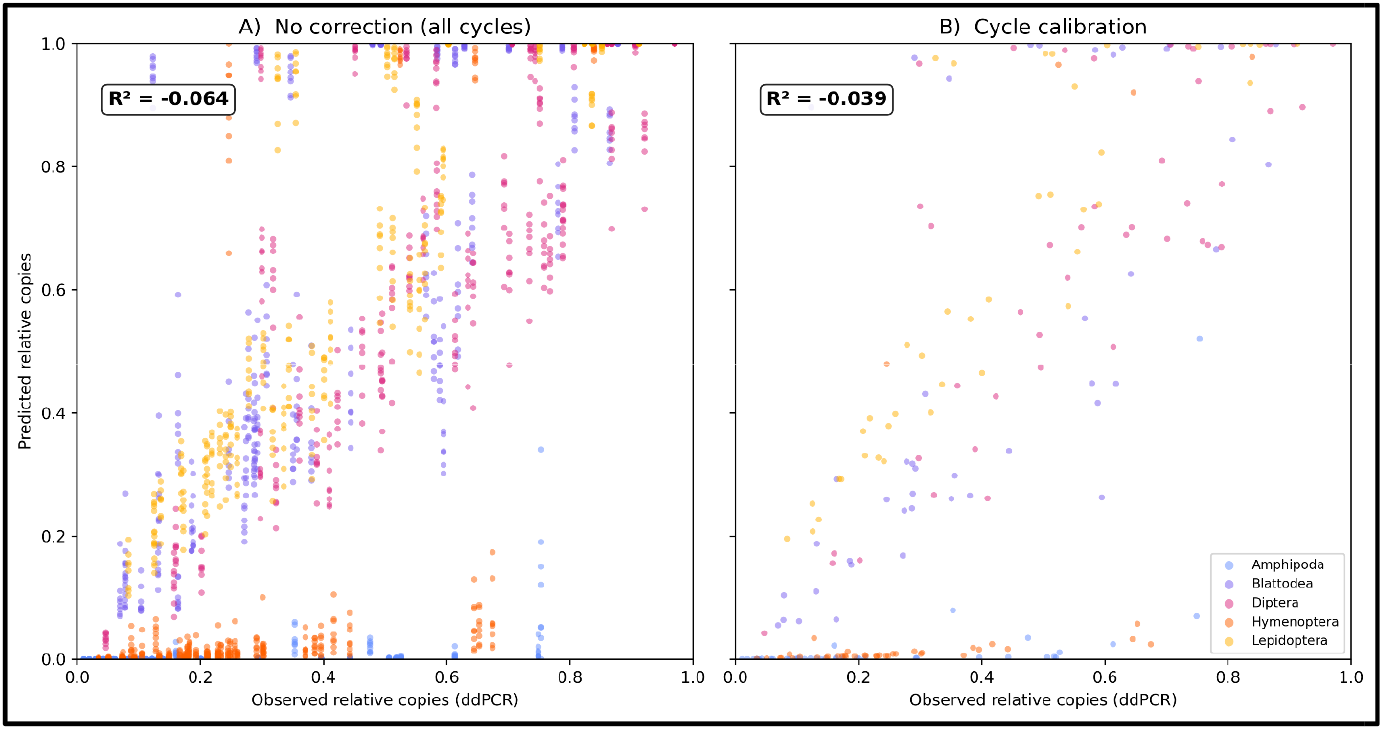
Observed relative mtDNA copy proportions (dPCR) vs. predicted proportions for all 81 mock communities, using raw metabarcoding reads without correction **(A)** and cycle-calibrated reads **(B).** Colors indicate order. R^2^ denotes the coefficient of determination as defined in Eq. 5.

### Mathematical framework for amplification bias

To formalize how this early fixation of primer-binding sites constrains amplification bias to the first two cycles of PCR, we defined a 2-stage system to describe the amplification process of a specific DNA template. Let ***N***_**0**_ be the number of starting templates, ***M*** be the number of amplicons that have been amplified with the forward primer only (medium-sized fragments), and ***S*** (short fragments) be the number of amplicons that have been amplified with both the forward and the reverse primer. Additionally, let ***p***_**1**_ be the probability that any ***N***_**0**_ successfully produces an ***M***, and let ***p***_**2**_ be the probability that any ***M*** produces an ***S***. Although the number of amplicons in each cycle strictly adheres to a binomial distribution, for the purposes of this model, we consider ***M*** and ***S*** as the expected fragment counts. After the primers have been replaced, the number of ***S*** simply doubles with each additional PCR cycle ***c***. The number of ***M*** after ***c*** cycles can then be described as:

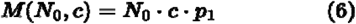

The quantity ***S*** after ***c*** cycles can be defined recursively as the sum of all ***S*** generated from ***M*** in the preceding cycle and the doubling of all ***S*** from the preceding cycle:

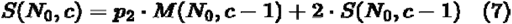

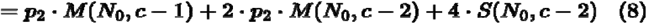

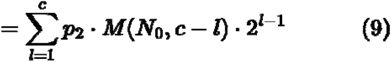

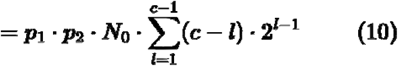

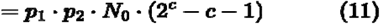

The full derivation is provided in supporting information 17. Since the bias of ***p***_**1**_ and ***p***_**2**_ cannot be independently observed, we define ***E*** as the amplification efficiency of a specific DNA target template where ***E*** = ***p***_**1**_ **· *p***_**2**_ which yields the final expression:

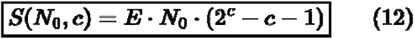

As we cannot observe the absolute number of ***S*** after sequencing, we define the relative reads ***R***_***r***_ of a given taxon ***i*** in a sample with ***j*** distinct taxa as:

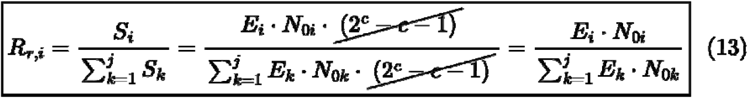

When utilizing relative reads, the cycle number is eliminated from the equation. By solving this for ***E***_***i***_ the relative Efficiency for a taxon ***i*** can be calculated as:

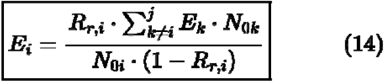

We selected *B. dubia* (Blattodea) as the reference taxon, against which we express ***E***_***i***_ values for all other taxa by setting ***R***_**ref**_ **= 1** (Table 1). For all 14 communities containing only the reference taxon and one other taxon (***j* = 2**), the general expression for ***E***_***i***_ simplifies, as the sum over ***k* ≠ *i*** collapses to the single reference term:

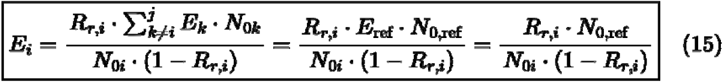

These relative ***E*** values were subsequently employed to predict the relative copy number from the relative reads measured after sequencing. The predicted relative copies for any taxon ***i*** can then be derived from the relative reads and efficiency values:

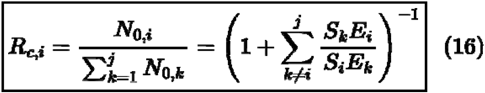

### mtDNA copy number calculation under different model assumptions

The mean efficiency values were calculated for all 14 two-species communities containing Blattodea (Mock community strategy) for the two models presented (cycle-dependent bias, equation 1; constant bias, equation 12; Table 2) and subsequently applied to predict mtDNA copies under both model assumptions (Figure 4 A, B).

**Table 2.**
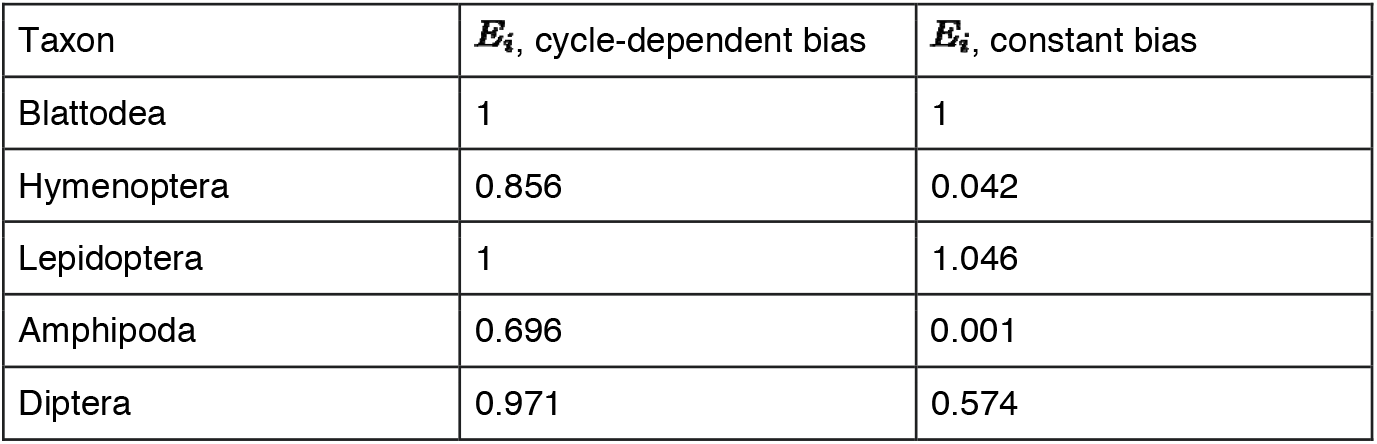
Mean relative amplification efficiencies of the 14 two-species calibration samples for the cycle-dependent bias model and the constant bias model.

| Taxon | $E_i$, cycle-dependent bias | $E_i$, constant bias |
| --- | --- | --- |
| Blattodea | 1 | 1 |
| Hymenoptera | 0.856 | 0.042 |
| Lepidoptera | 1 | 1.046 |
| Amphipoda | 0.696 | 0.001 |
| Diptera | 0.971 | 0.574 |

**Figure 4.**
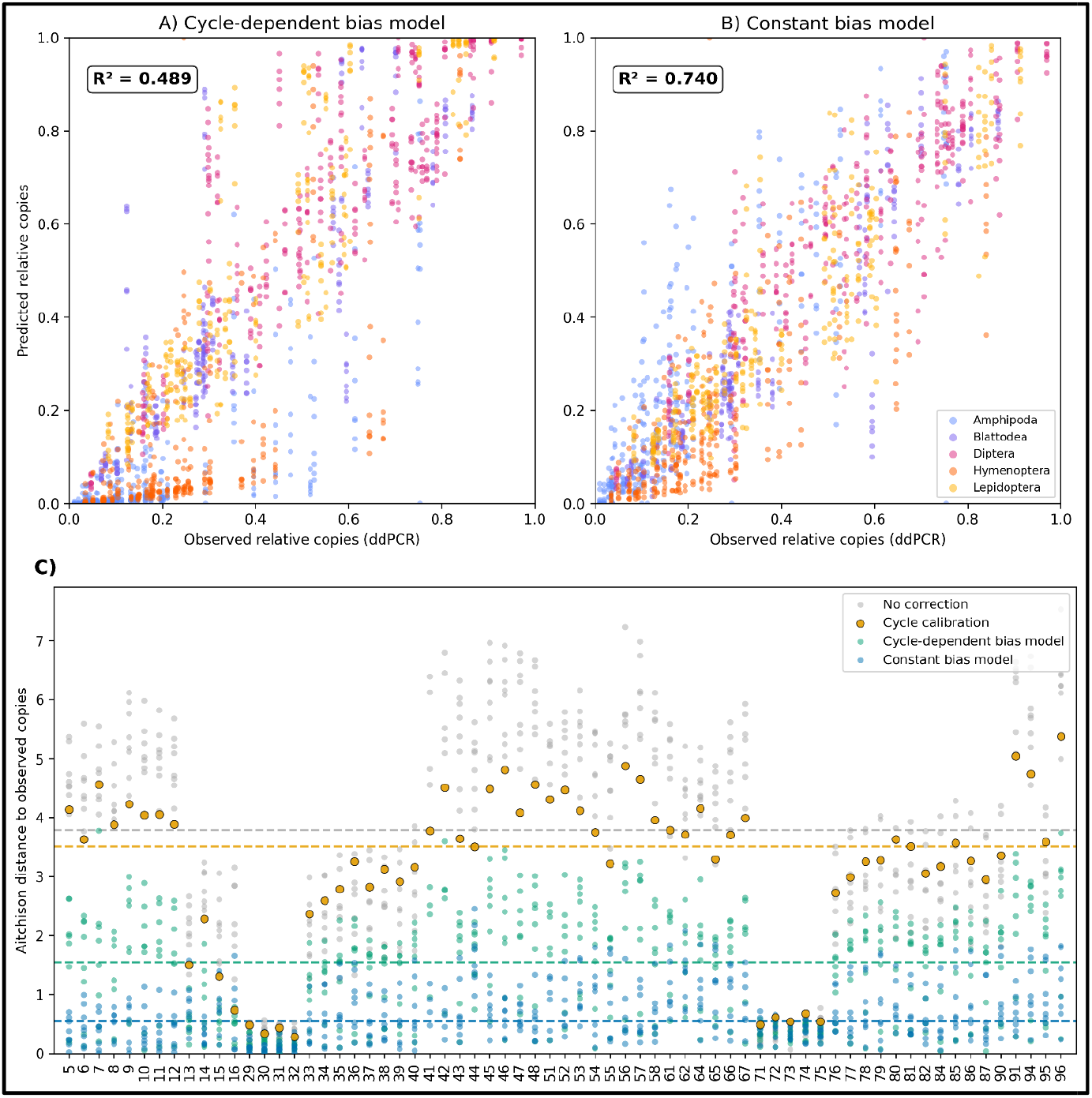
Predictive accuracy of three approaches for correcting PCR amplification bias in metabarcoding data. **(A)** Predicted versus observed relative copy numbers (dPCR) for the cycle-dependent bias model. **(B)** Predicted versus observed relative copy numbers (dPCR) for the constant bias model. Predictions are made for the 67 communities that were not used for calibration. R^2^ denotes the coefficient of determination as defined in Eq. 5. **(C)** Aitchison distance between predicted and observed (dPCR) taxon proportions for each of the 67 communities across all PCR cycle counts tested (c = 6–20). Dashed horizontal lines indicate the median Aitchison distance per approach. Samples containing 0 reads are not shown.

Both models improved the prediction of mtDNA copy numbers compared to no correction, with the constant bias model presented in this study providing the better prediction (cycle-dependent bias model R^2^ = 0.489, constant bias model R^2^ = 0.740). Since both models are calibrated against the relative reads after cycle 20, they provide near-identical predictions at c=20 by construction. The constant bias model performed better at predicting mtDNA copy numbers for the cycle numbers 6-16, while the difference is minor for c=18 (supporting information 18, supporting information 19). The constant bias model performed better across all taxonomic orders, with the difference being most pronounced for the severely biased Amphipoda and Hymenoptera (Supporting information 20).

The median Aitchison distance between predicted and observed taxon proportions was highest for uncorrected reads (3.78) and the cycle calibration method (3.51), indicating that the cycle calibration method does not substantially reduce compositional bias at the community level. The cycle-dependent bias model reduced the median distance to 1.55, while the constant bias model provided the strongest improvement, reducing it to 0.54. Samples 29–32 (Blattodea + Lepidoptera) and 71–75 (Blattodea, Lepidoptera, and Diptera) showed consistently low Aitchison distances across all methods, including no correction. This is expected given that the relative amplification efficiencies of these taxa are closest to 1 (Table 2), meaning their read proportions already closely reflect mtDNA copy proportions regardless of the correction applied (Figure 4C).

Mean relative amplification efficiencies for the constant bias model were also calculated for the BF3 primer pair. Compared to the fwh2 primer pair, Amphipoda was overrepresented (***E***_***i***_ **= 1.30**) and Diptera showed nearly unbiased amplification (***E***_***i***_ **= 0.97**) in the BF3 samples, while Hymenoptera remained strongly underrepresented with both primer pairs (supporting information 21). The constant bias model also improved the predicted mtDNA copy numbers for the BF3 primer pair (R^2^ = 0.43 for the uncorrected relative reads and R^2^ = 0.90 for the corrected reads, supporting information 22).

## Discussion

Our replicated mock community study showed that PCR amplification bias in metabarcoding is taxon-specific and non-exponential. The compositional nature of sequencing data further complicates the direct assessment of correction factors, as the demonstrated taxon-specificity of amplification bias does not invariably result in a taxon being consistently under-or overrepresented. Rather, the taxon-specific signal is masked by the compositional characteristics of the metabarcoding data.

In contradiction to hypothesis 1, species biomass proportions are not well reflected in relative mtDNA copies. The relative rank of species’ biomass was only congruent with mtDNA copy rank for half of the data points. The fold-change of relative sample share was out of the 1.5-fold range in about half of the cases and revealed systematic underrepresentation especially for Amphipoda, as well as overrepresentation for Diptera. In this context it needs to be noted, that compositional data is bounded at 100%. Therefore, underrepresentation can drive the fold-change deviation arbitrarily high, while overrepresentation is inherently capped – making fold-change asymmetric across co-occurring taxa. Furthermore, the selected threshold of a 1.5-fold change is arbitrary, and technically, any bias in fold-change representation distorts the quantitative quality of the data. The discrepancy between relative biomass and relative mtDNA copy numbers confirms the observations of several studies that report differences in mtDNA copy number of humans and other mammals depending on several influence factors like tissue type, individual and age (Naue et al., 2024; Rath et al., 2024; Wachsmuth et al., 2016).

Regarding hypothesis 2, which stated that metabarcoding read numbers do not reflect mtDNA copy numbers without correction, we observed two different scenarios: when the differences in amplification efficiency are minor, like for Blattodea, Diptera, and Lepidoptera in our dataset, mtDNA copy numbers are reflected in metabarcoding reads. This is confirmed by a high rank concordance for communities only containing these taxa as well as by low fold-change values between mtDNA copies and metabarcoding reads. When the amplification efficiency of any taxon in a community is low, the representation of mtDNA copies in metabarcoding reads is highly biased, leading to deviations of as much as 12-fold underrepresentation for Amphipoda in this dataset. We therefore reject hypothesis 2 for communities that only contain taxa with minor amplification efficiency differences, whose reads do reflect mtDNA copy numbers, but confirm it for communities that contain taxa such as Amphipoda and Hymenoptera, for which strong taxon-specific bias decouples metabarcoding reads from mtDNA copy numbers.

Integrating our findings from testing hypothesis 1 and 2, we show that from input biomass to metabarcoding biases are accumulated. Consequently, we observed low rank concordances in most of the taxa, as well as high values of underrepresentation for the taxa with low amplification efficiencies Amphipoda and Hymenoptera. Given the fact that it is very likely for mixed samples to contain taxa that amplify poorly and therefore distort the accuracy of representation in reads, a correction is needed to enable an accurate representation of the sample’s community. Therefore, and in line with previous work (Saitoh et al., 2016; Thomas et al., 2016), we see the necessity for a correction method that enables the inference of the original template mtDNA copy number of a sample from metabarcoding read data.

Opposite to our initial expectations from hypothesis 3, PCR cycle calibration did not offer a sufficient correction strategy, as the data did not show systematic changes in read composition with increasing cycle numbers. Thus, the cycle calibration strategy did not predict mtDNA copy numbers any better than just using uncorrected reads themselves. Given this result, we consequently turned to the mock community strategy as suggested by (Saitoh et al., 2016; Thomas et al., 2016). Under the assumption of an exponentially compounding amplification bias, as described by Eq. 1, both methods should yield similar efficiency estimates. Our empirical data showed that, however, the mock community strategy predicted mtDNA copy numbers considerably better than cycle calibration, although it performed poorly for communities containing poorly amplifying taxa. It should be noted that any model correctly predicting the direction of bias will outperform uncorrected reads; a meaningful test of the underlying mechanism therefore requires evaluating model predictions at cycle numbers different from those used for calibration. When applying this test, our data showed that the mock community strategy’s performance deteriorated with decreasing cycle numbers.

Although our data do not support the cycle calibration strategy suggested by Silverman et al. (Silverman et al., 2021), they corroborate another finding of that work: bias-inducing primer mismatches are replaced by perfectly matching primer sequences after the first few PCR cycles, a mechanism also described by Wu et al. (Wu et al., 2009). Silverman et al. (Silverman et al., 2021), however, still assume an exponential nature of PCR amplification bias, attributing it instead to non-primer-mismatch sources such as a sequence’s GC content, its propensity for secondary structure formation, or sequence length. Our mock community data, however, do not support an exponential nature of this process, in contrast to the mechanisms proposed. Moreover, the differing efficiencies we estimated for the second primer pair, BF3, make primer mismatch a considerably more likely explanation for the observed bias than these alternative mechanisms. Taken together, our data therefore support the primer replacement mechanism rather than an exponential, primer-independent bias model. We therefore defined taxon-specific amplification efficiency as the probability with which forward and reverse primers each produce a copy of the original template — a probability governed by primer-template mismatches. Because primer binding sites are replaced by actual primer sequences within the first two PCR cycles, bias does not compound with increasing cycle numbers, consistent with the experimental findings of Krehenwinkel et al. (Krehenwinkel et al., 2017).

Through our calculation of a non-exponential species-specific amplification efficiency relative to the amplification efficiency of a reference species in the sample, the inference of input mtDNA copy numbers in a sample is possible with high accuracy. Relative amplification efficiencies were applicable independent of the species composition of the sample and the number of PCR cycles used for amplification. The correction increased the accuracy of the mtDNA copy estimation in all cases, especially in the fwh2 primer pair, for which the metabarcoding reads were unsuitable for predicting mtDNA copy number without correction in most cases. For the BF3 primer pair, which provided slightly better mtDNA copy number representation in metabarcoding reads, the accuracy-increasing effect of the correction was only relevant in one species, while the spread of error was reduced for all species. Thus, overall, we observe a beneficial effect of corrections especially for primer pairs that do not provide accurate representation of mtDNA copies through metabarcoding reads. These findings about PCR amplification bias are relevant for our fundamental understanding about the mechanistic processes of PCR and the possibility to derive quantitative information. It remains to be tested whether the constant-bias correction model generalizes to other mitochondrial markers commonly used in metabarcoding, such as cytochrome b, 16S rDNA or 12S rDNA.

However, the practical applications are currently limited by several factors: Due to the high variation of mtDNA copies per input biomass, inferring a precise biomass estimate from measured or calculated copy numbers does not seem feasible. The variation of chloroplast DNA in different plant species and tissues as observed by Kamenova et al (Kamenova et al., 2025), indicates that this problem is likely inherent to many metabarcoding applications, including animal and plant metabarcoding. Apart from adding a severe bias to biomass estimation from mtDNA in bulk samples, this also presents a potential problem regarding biomass estimation from environmental DNA. The estimation of mtDNA copy number in eDNA samples is further complicated not only by the large variation of mtDNA content in tissues, but also by potential differences in shedding rates from different taxa and life stages. Therefore, we consider mtDNA copy number variation to be a major hurdle in mtDNA-based biomass quantification: Even with the ability to precisely infer mtDNA copy numbers, the estimation of animal biomass and more so, abundance, based on DNA sequencing information is highly inaccurate at the current state of research. With the level of variation of biomass in individuals and mtDNA copy number in tissues and life stages, a substantial amount of further research is needed to overcome the limitation of DNA-based quantification to the level of (mt)DNA copy numbers. A potential alternative lies in the use of nuclear single copy markers that show substantially less copy number variation. However, current barcoding identification methods for animals are focused on mitochondrial markers and therefore offer the most comprehensive databases. The use of tissue- and DNA extract-based spike ins has been heavily discussed in the context of quantitative metabarcoding (Iwaszkiewicz-Eggebrecht et al., 2026; Luo et al., 2023; Tkacz et al., 2018) and could bear potential to function as an amplification reference added to biological samples.

Still, the inference of relative mtDNA copy numbers is only accurate if all species contained in the sample and their respective relative amplification efficiencies are known. Even though the proportional relationships among the known target species remain constant when not all sample contents are known, the absolute read proportions become highly skewed in this case. The prerequisite of knowing all species and their relative amplification efficiencies in a sample can probably never be met for complex samples of highly diverse taxa like insects in Malaise trap samples, or samples that contain many non-target templates such as eDNA samples. It is, however, thinkable that the method can be applied to samples with limited taxa scope, e.g. freshwater macroinvertebrate samples acquired for ecological assessment within the European Water Framework Directive (Hering et al., 2018). Additionally, to infer mtDNA copy numbers, the amplification efficiency of the target species in relation to a reference species that is also contained in the sample must be calculated. For this purpose, a two-species mock community must be quantified for all taxa in question, independently from the ecological sample of interest. For the method to become feasible in practical applications there is thus a need to find a faster and cheaper way to determine relative amplification efficiencies. It is currently an open question to what extent it is realistic to determine general amplification efficiencies that are applicable to species, genera, or even families relative to e.g. a fixed spike-in as a reference, especially for markers like COI and phylogenetically diverse groups such as insects.

It is important to keep in mind that all these approaches would only suffice to calculate accurate mtDNA copy number and not biomass or abundance for the reasons stated above. We therefore believe that this study contributes to understanding the mechanistic details of PCR amplification but that a substantial amount of further research is required to achieve practical applicability of our findings.

## Supporting information

Overview of the supporting information

## Author contributions

DB, LW, and FL designed the study concept. LW and DB conducted the laboratory work. DB and LW conducted the bioinformatic processing and data analysis. DB and KK conceptualized the mathematical description of the mechanism. LW wrote the first draft of the manuscript, which was revised by all coauthors.

## Acknowledgement and funding

We would like to thank the Dr. Herzog-Sellenberg-Foundation (Stifterverband für die Deutsche Wissenschaft) for funding this study. Furthermore, this study was supported by Collaborative Research Center (CRC) RESIST funded by the Deutsche Forschungsgemeinschaft (DFG, German Research Foundation) – CRC 1439/2 – project number: 426547801. We thank Julian Enß and Franziska Wenskus for contributing study organisms. We thank Kamilla Adam, Marie Borowski, and Elena Kuhn for assistance with sample processing in the lab. We thank Arne J. Beermann, Iris Madge-Pimentel, and Ryan P. Kelly for feedback on the study and Arne Beermann for comments on the manuscript. Open Access funding enabled and organized by Projekt DEAL.

## Data accessibility statement

Demultiplexed raw read data for this study have been uploaded to the European Nucleotide Archive under the accession number PRJEB110937.

## Ethical statement

All specimens were invertebrates from breeding cultures, not wild-collected. No ethical approval or permits were required, as invertebrates are not subject to German or EU animal welfare legislation.

