## Supplementary figures and images for "Characterizing mitochondrial copy number variation and PCR amplification bias as sources of quantitative constraints in DNA metabarcoding"

### Supporting information 4

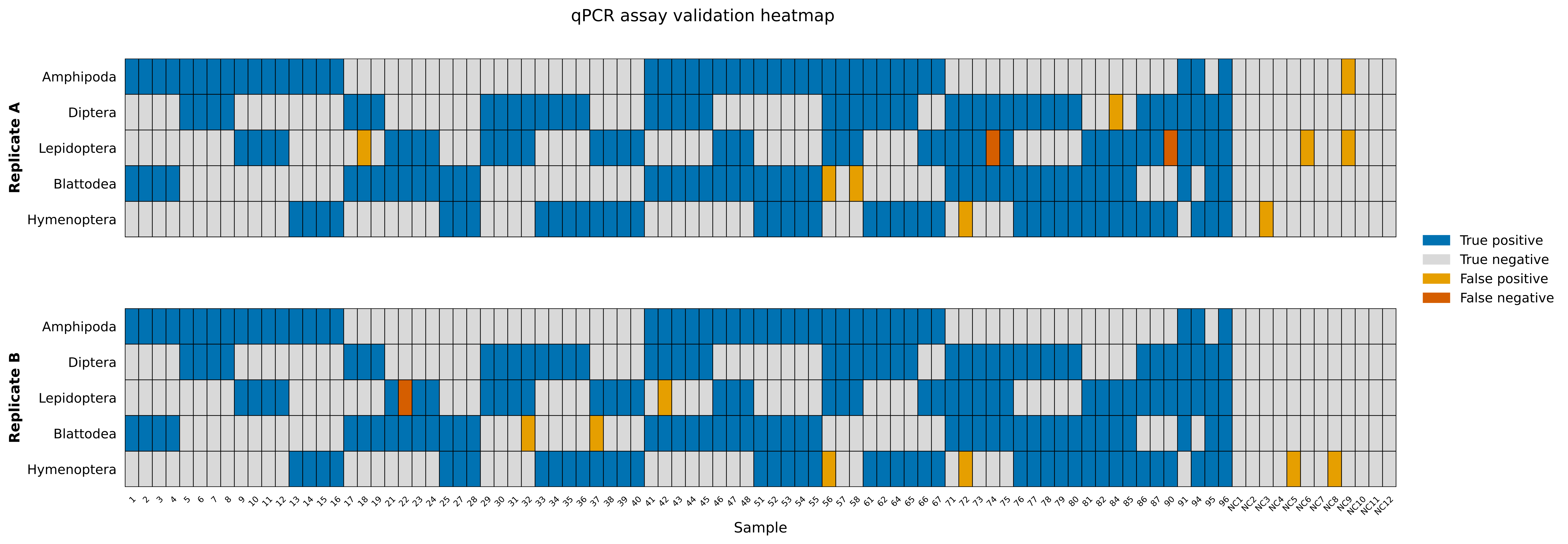

### Supporting information 7

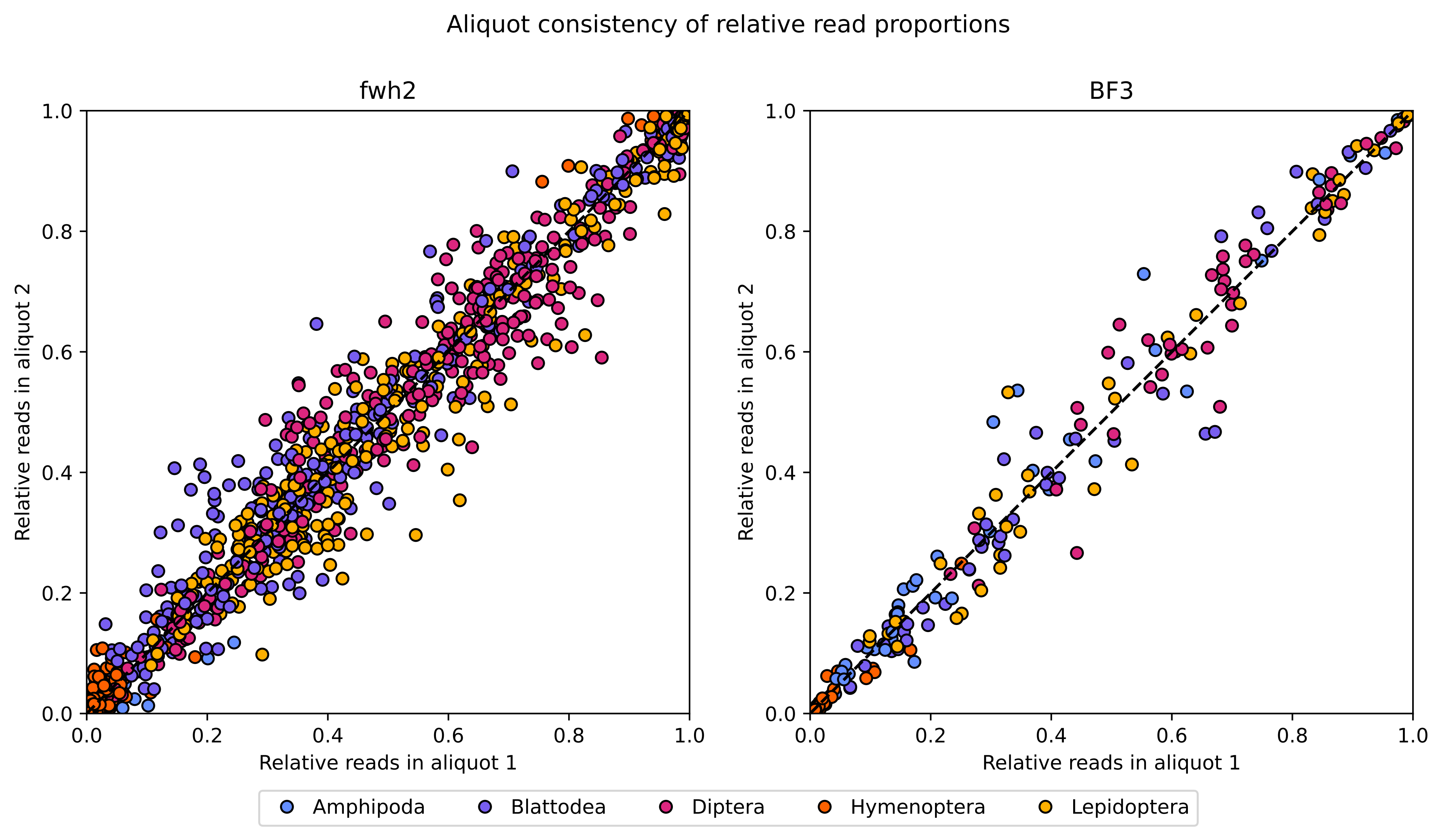

### Supporting information 8

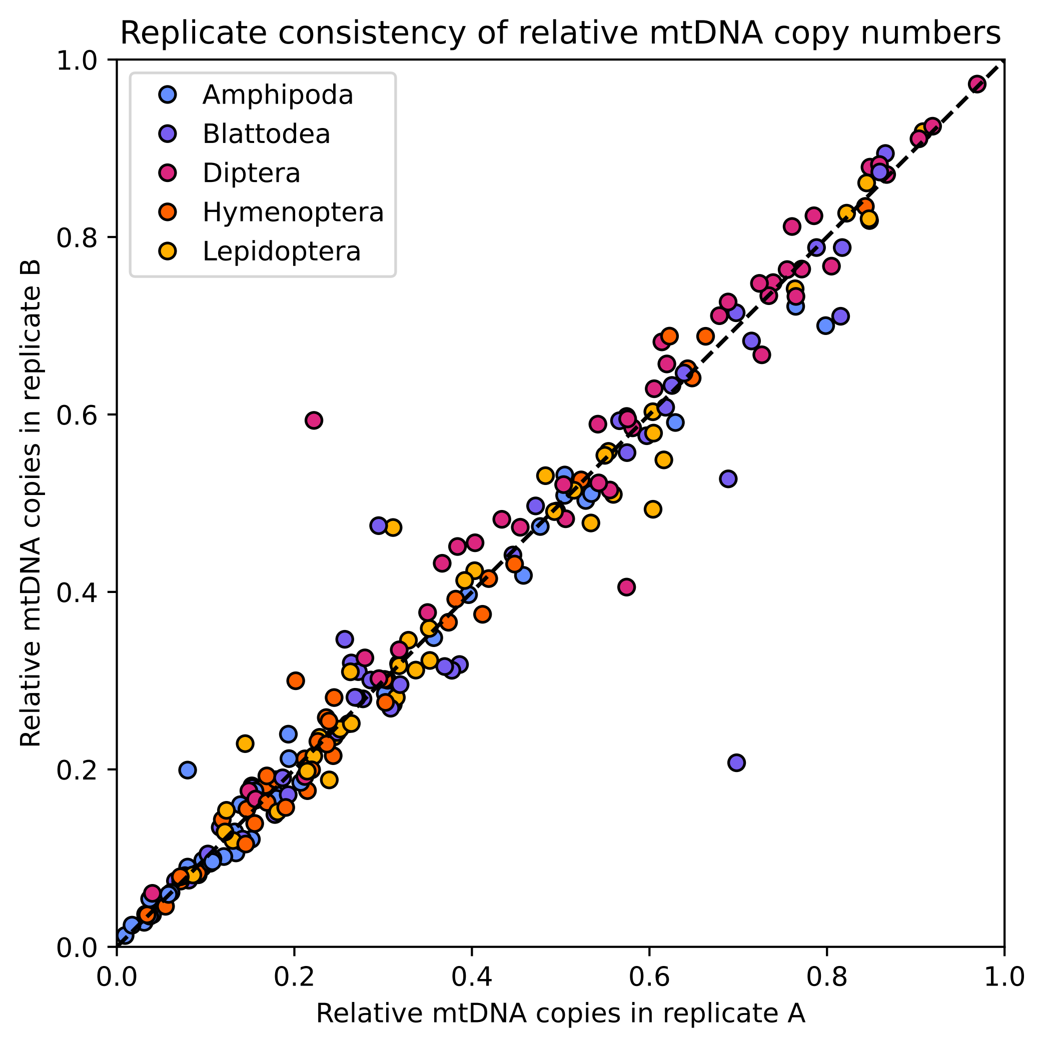

### Supporting information 12

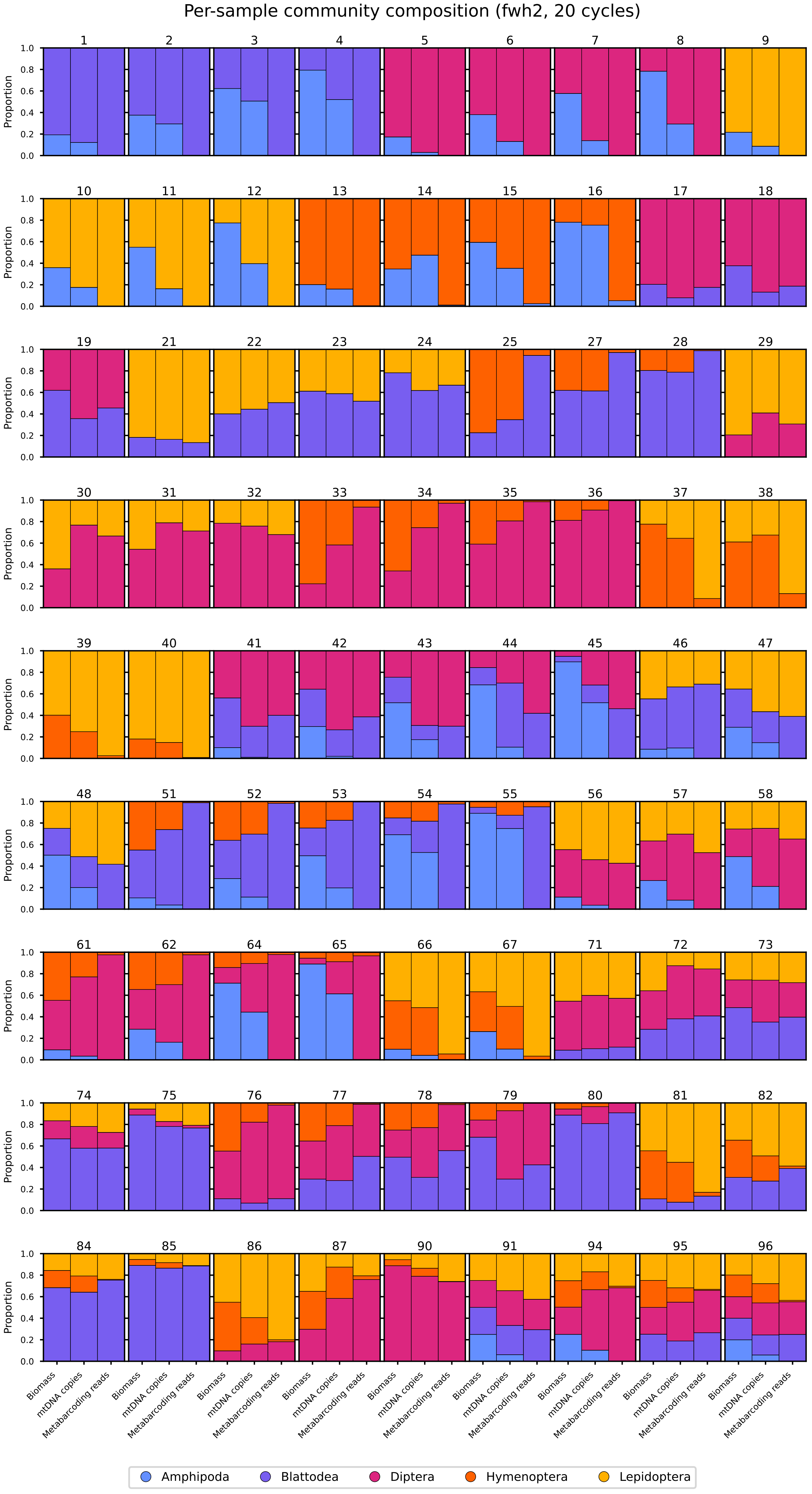

### Supporting information 13

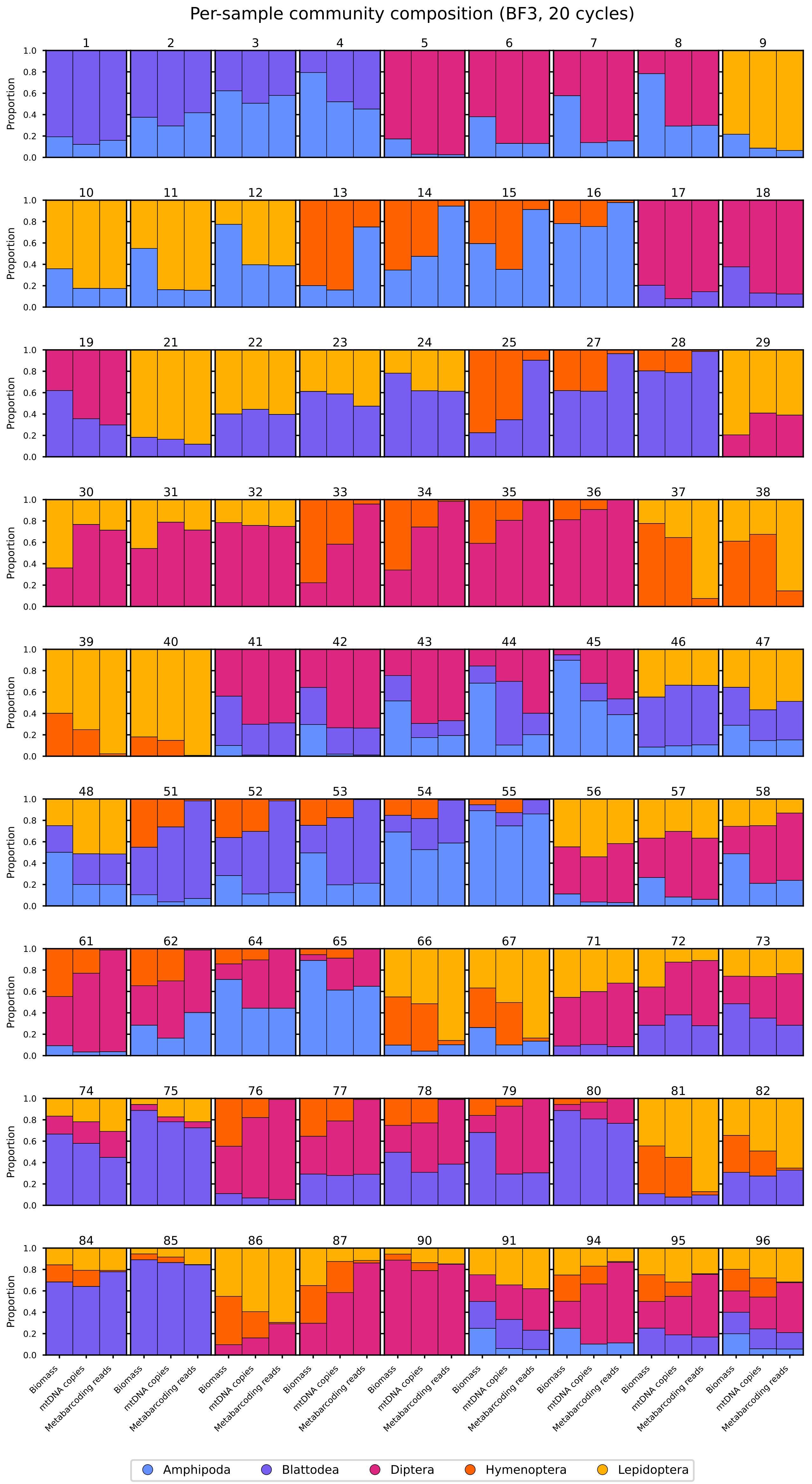

### Supporting information 14

Relative metabarcoding read proportions across PCR cycles (fwh2)

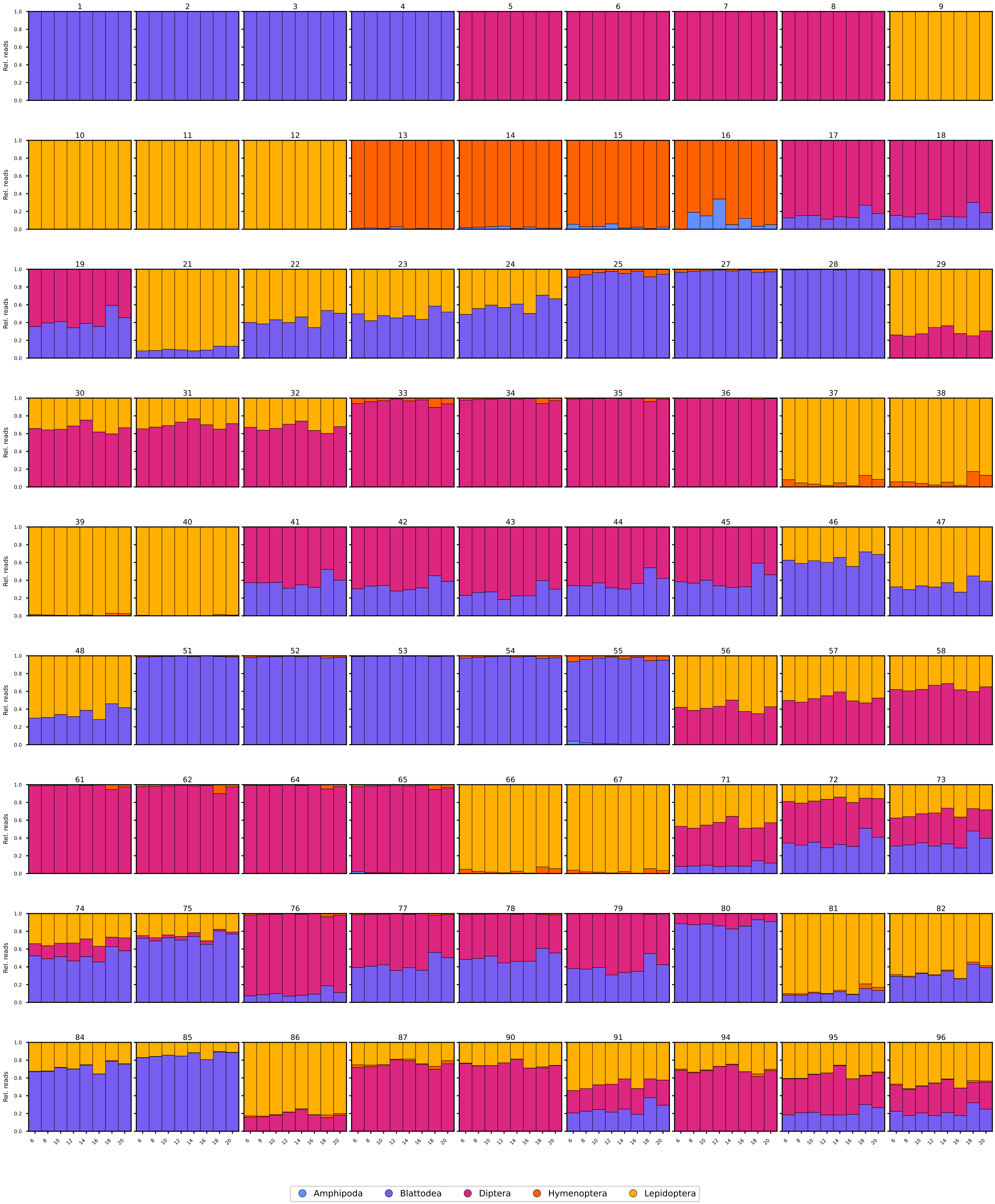

### Supporting information 18

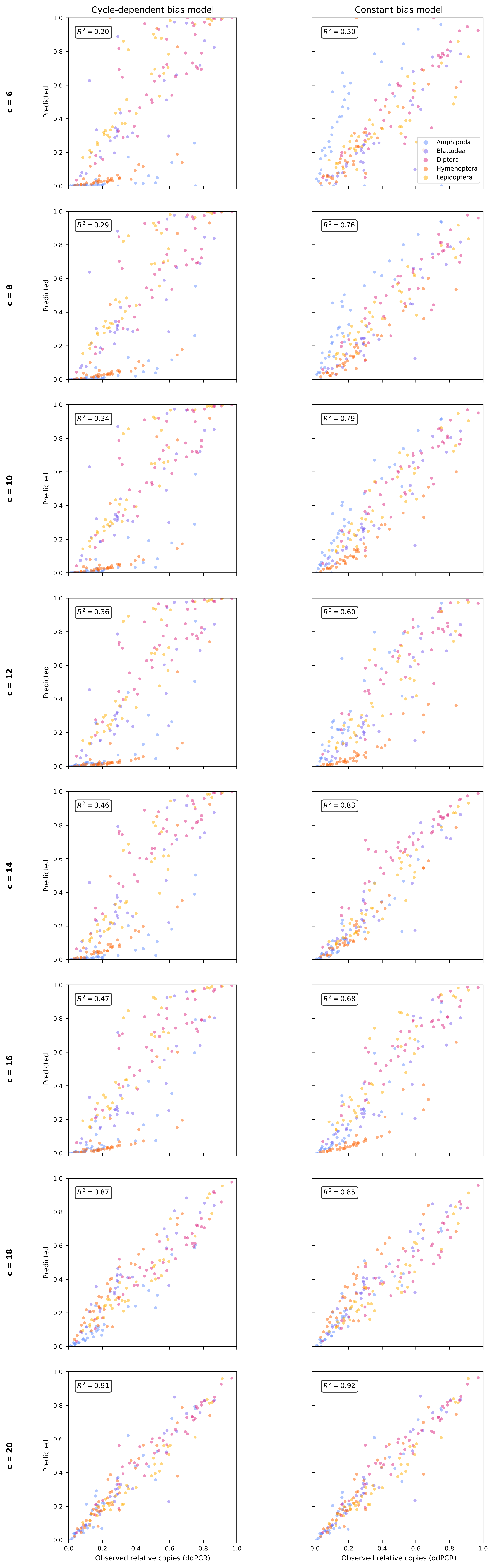

### Supporting information 20

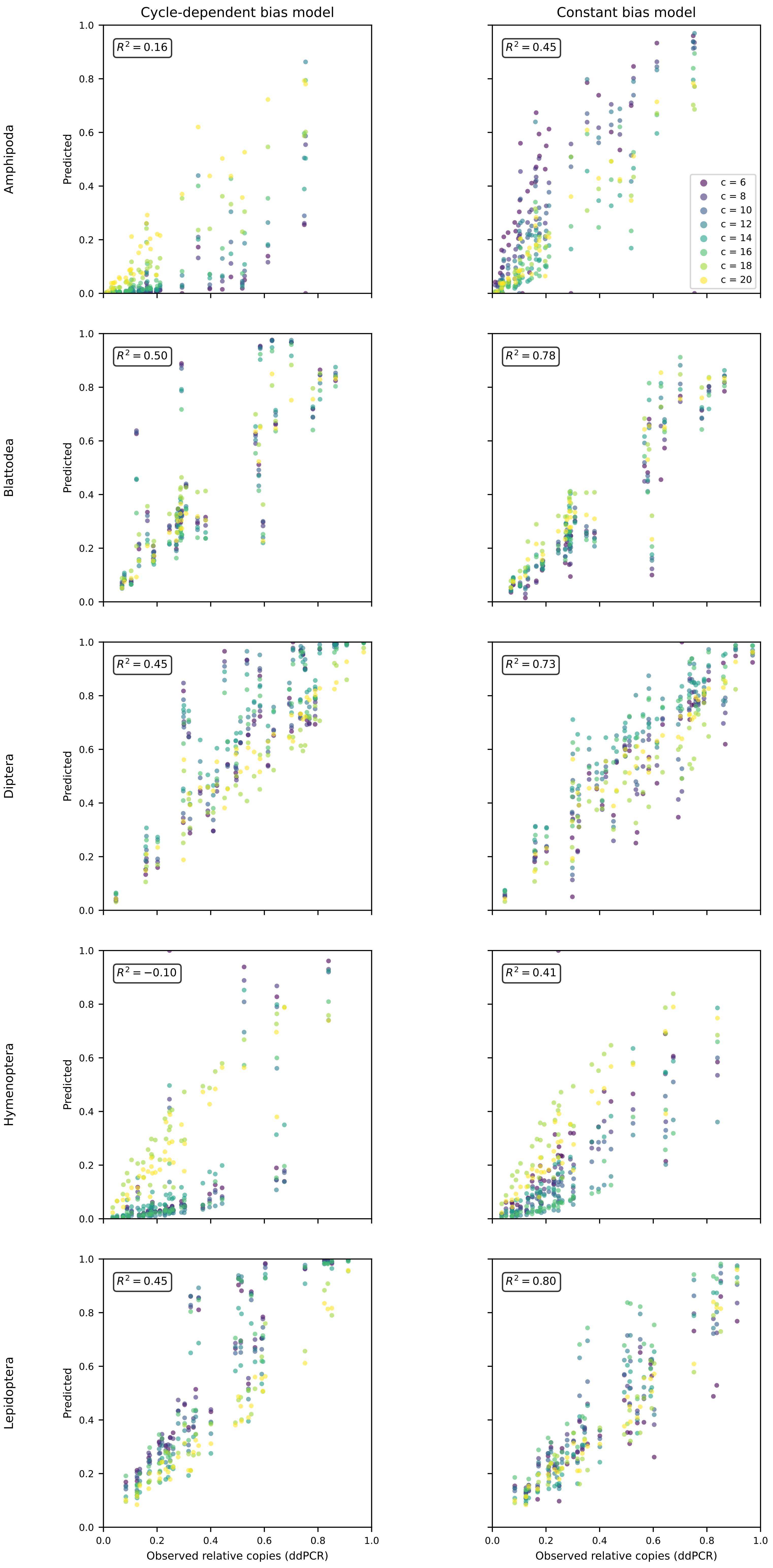

### Supporting information 22

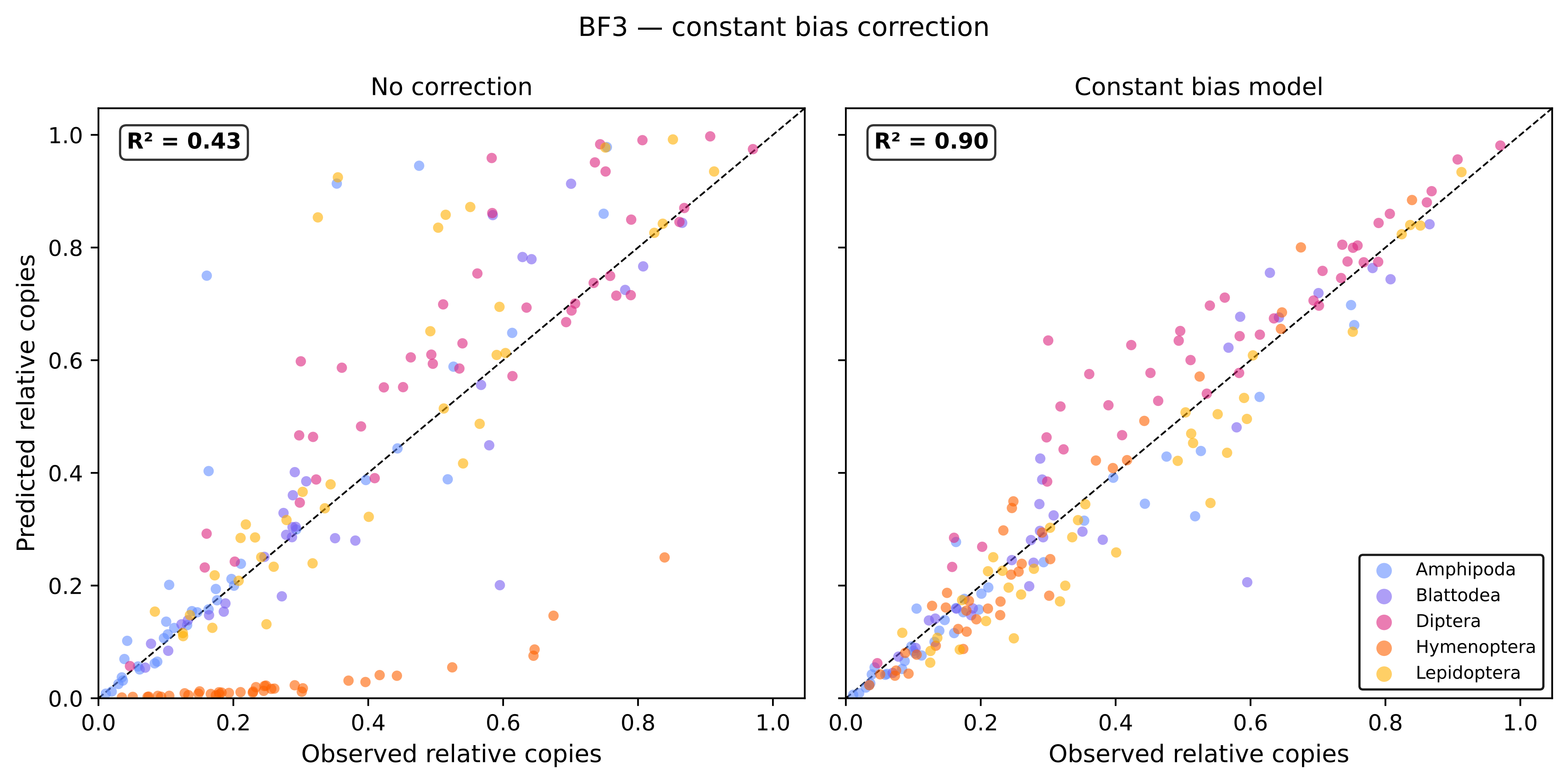
