## Supporting information 10 for "Characterizing mitochondrial copy number variation and PCR amplification bias as sources of quantitative constraints in DNA metabarcoding": supporting information 10.py

```
import numpy as np
import pandas as pd
from pathlib import Path
import matplotlib.pyplot as plt
import seaborn as sns
from matplotlib.colors import ListedColormap, BoundaryNorm
import matplotlib.patches as mpatches
from scipy.stats import linregress, wilcoxon

ORDER_COLORS = ["#648FFF", "#785EF0", "#DC267F", "#FE6100", "#FFB000"]
ORDER_NAMES = ["Amphipoda", "Blattodea", "Diptera", "Hymenoptera", "Lepidoptera"]
ORDER_PALETTE = dict(zip(ORDER_NAMES, ORDER_COLORS))


def supporting_information_4(supporting_information_3: Path) -> None:
    """Create the qPCR validation heatmap (Supporting Information 4).

    Before running metabarcoding, we checked with taxon-specific qPCR assays
    whether each target insect order was actually detectable in each community
    sample. This function reads those qPCR results and produces a colour-coded
    grid (heatmap) where rows are insect orders, columns are samples, and each
    cell is coloured by detection outcome: true positive (detected as expected),
    true negative (absent as expected), false positive (detected but absent),
    or false negative (absent but expected). Both extraction replicates (A and B)
    are shown as separate panels. The result is saved as Supporting Information 4.
    """
    # --- Data loading ---
    supporting_information_3 = pd.read_excel(supporting_information_3)

    # Fixed display order of the five qPCR target taxa (rows of the heatmap)
    TARGET_ORDER = ["Amphipoda", "Diptera", "Lepidoptera", "Blattodea", "Hymenoptera"]

    # Sample order follows the original table; negative controls are already last
    SAMPLE_ORDER = supporting_information_3["sample"].unique().tolist()

    # --- Colour mapping for the four detection states ---
    STATE_LABELS = {
        0: "True positive",
        1: "True negative",
        2: "False positive",
        3: "False negative",
    }

    # Colorblind-friendly palette based on Okabe-Ito
    STATE_COLORS = {
        0: "#0072B2",  # TP  – blue
        1: "#D9D9D9",  # TN  – light gray
        2: "#E69F00",  # FP  – orange
        3: "#D55E00",  # FN  – vermillion
    }

    # Discrete colormap: map integer states 0–3 to the four colours above
    cmap = ListedColormap([STATE_COLORS[i] for i in range(4)])
    # Boundaries centred around each integer so imshow bins values correctly
    norm = BoundaryNorm([-0.5, 0.5, 1.5, 2.5, 3.5], cmap.N)

    # --- Helper: build a (target × sample) matrix for one replicate ---
    def make_matrix(replicate):
        sub = supporting_information_3[
            supporting_information_3["replicate"] == replicate
        ]
        pivot = sub.pivot(index="Target Name", columns="sample", values="state")
        # Enforce consistent row/column order across replicates
        pivot = pivot.reindex(index=TARGET_ORDER, columns=SAMPLE_ORDER)
        return pivot

    mat_a = make_matrix("A")
    mat_b = make_matrix("B")

    # --- Figure setup: two vertically stacked heatmaps (one per replicate) ---
    fig, axes = plt.subplots(
        2, 1, figsize=(18, 6.5), sharex=True, gridspec_kw={"hspace": 0.4}
    )

    for ax, mat, label in zip(axes, [mat_a, mat_b], ["Replicate A", "Replicate B"]):
        # Draw the heatmap for this replicate
        ax.imshow(
            mat.values, aspect="auto", cmap=cmap, norm=norm, interpolation="nearest"
        )

        # Y-axis: target taxa labels
        ax.set_yticks(range(len(TARGET_ORDER)))
        ax.set_yticklabels(TARGET_ORDER, fontsize=11)
        ax.set_ylabel(label, fontsize=12, fontweight="bold")

        # X-axis: one tick per sample (labels only on the bottom panel)
        ax.set_xticks(range(len(SAMPLE_ORDER)))

        # Remove default tick marks and axis spines for a cleaner look
        ax.tick_params(left=False, bottom=False)
        for spine in ax.spines.values():
            spine.set_visible(False)

        # Draw a grid between cells using minor ticks placed at cell boundaries
        ax.set_xticks([x - 0.5 for x in range(1, len(SAMPLE_ORDER))], minor=True)
        ax.set_yticks([y - 0.5 for y in range(1, len(TARGET_ORDER))], minor=True)
        ax.grid(which="minor", color="black", linewidth=0.6)
        ax.tick_params(which="minor", length=0)

        # Draw an explicit outer border around the entire matrix
        ax.set_xlim(-0.5, len(SAMPLE_ORDER) - 0.5)
        ax.set_ylim(len(TARGET_ORDER) - 0.5, -0.5)
        for x in [-0.5, len(SAMPLE_ORDER) - 0.5]:
            ax.axvline(x, color="black", linewidth=0.8)
        for y in [-0.5, len(TARGET_ORDER) - 0.5]:
            ax.axhline(y, color="black", linewidth=0.8)

    # X-axis labels only on the bottom panel, rotated for readability
    axes[-1].set_xticklabels(
        SAMPLE_ORDER, rotation=45, ha="right", rotation_mode="anchor", fontsize=7
    )
    axes[-1].set_xlabel("Sample", fontsize=12)

    fig.suptitle("qPCR assay validation heatmap", fontsize=14, y=0.97)

    # --- Legend (placed to the right of the heatmaps) ---
    legend_patches = [
        mpatches.Patch(color=STATE_COLORS[i], label=STATE_LABELS[i]) for i in range(4)
    ]
    fig.legend(
        handles=legend_patches,
        loc="center left",
        bbox_to_anchor=(0.96, 0.5),
        frameon=False,
        fontsize=11,
    )

    # Leave 5 % right margin for the legend; avoid tight_layout (incompatible
    # with sharex and manual legend placement). bbox_inches="tight" in savefig
    # handles the final bounding box.
    fig.subplots_adjust(right=0.95)

    # --- Export ---
    plt.savefig("supporting information 4.pdf", dpi=300, bbox_inches="tight")


def aggregate_metabarcoding_data(
    supporting_information_1: Path, supporting_information_6: Path
) -> pd.DataFrame:
    """Load and clean the raw metabarcoding sequencing data (Supporting Information 6).

    Metabarcoding produces a table of DNA sequence counts ("reads") for each
    taxon detected in each sample. This function loads that raw table, sums
    reads across sequencing lanes for each sample/taxon/primer/PCR-cycle
    combination, and then removes any reads belonging to taxa that were not
    intentionally added to that community (co-amplified non-target sequences).
    Three samples (26, 92, 93) that were not part of the controlled experiment
    are also excluded. Combinations where a taxon was expected but produced
    zero reads are explicitly recorded as zero rather than left as missing,
    so that all downstream calculations see complete data.
    Returns the cleaned read table.
    """
    # --- Load and aggregate raw read data ---
    supporting_information_6 = pd.read_parquet(supporting_information_6)

    supporting_information_6 = (
        supporting_information_6.groupby(
            by=["sample", "aliquot", "cycles", "order", "primer"]
        )
        .sum()["read_count"]
        .reset_index()
    )

    # --- Load expected community composition ---
    supporting_information_1 = pd.read_excel(supporting_information_1)
    expected_taxa = supporting_information_1[["sample", "order"]].drop_duplicates()

    # --- Filter to samples present in SI 1 (drops e.g. 26, 92, 93) ---
    valid_samples = set(expected_taxa["sample"])
    supporting_information_6 = supporting_information_6[
        supporting_information_6["sample"].isin(valid_samples)
    ]

    # Total reads per primer before removing co-amplifications
    pre_filter_reads = supporting_information_6.groupby("primer")["read_count"].sum()

    # --- Build a complete design grid ---
    # All experimental conditions that were actually sequenced
    design = supporting_information_6[
        ["sample", "aliquot", "cycles", "primer"]
    ].drop_duplicates()

    # Cross-join each condition with the expected orders for that sample
    design_grid = design.merge(expected_taxa, on="sample", how="left")

    # --- Merge actual read counts onto the grid; fill missing with 0 ---
    result = design_grid.merge(
        supporting_information_6,
        on=["sample", "aliquot", "cycles", "order", "primer"],
        how="left",
    )
    result["read_count"] = result["read_count"].fillna(0).astype(int)

    # Total reads per primer after removing co-amplifications
    post_filter_reads = result.groupby("primer")["read_count"].sum()

    print("\n=== Metabarcoding read filtering (samples 26/92/93 excluded) ===")
    for primer in ["fwh2", "bf3"]:
        pre = pre_filter_reads[primer]
        post = post_filter_reads.get(primer, 0)
        removed = pre - post
        pct = removed / pre * 100 if pre > 0 else 0
        print(
            f"  {primer}:\n"
            f"    Before filtering: {pre:,}\n"
            f"    After filtering:  {post:,}\n"
            f"    Removed:          {removed:,} ({pct:.2f}%)"
        )

    # Sort by primer (fwh2 before bf3), then sample/cycles/aliquot ascending
    result = result.sort_values(
        by=["primer", "sample", "cycles", "aliquot"],
        ascending=[False, True, True, True],
    )

    return result


def absolute_reads_to_relative_reads(
    aggregated_read_data: pd.DataFrame,
) -> pd.DataFrame:
    """Convert raw read counts to relative proportions and verify extraction reproducibility.

    Because total sequencing depth varies between samples and runs, raw read
    counts are not directly comparable. This function divides each taxon's
    read count by the total reads in that sample/aliquot/cycle group, giving
    the relative proportion (0–1) of each taxon. It then checks reproducibility:
    each sample was extracted in two aliquots, and a Wilcoxon signed-rank test
    (a non-parametric pairwise test) checks whether the two aliquots give
    consistently similar proportions. After the check the two aliquots are
    pooled (summed) into a single value per sample.
    Returns the table with relative proportions added.
    """
    # --- Compute read sum per (sample, aliquot, cycles, primer) ---
    grouped_data = (
        aggregated_read_data.groupby(by=["sample", "aliquot", "cycles", "primer"])
        .sum()["read_count"]
        .reset_index()
        .rename(columns={"read_count": "read_sum"})
    )

    # Merge read_sum back so each row knows its group total
    aggregated_read_data = pd.merge(
        left=aggregated_read_data,
        right=grouped_data,
        how="left",
        on=["sample", "aliquot", "cycles", "primer"],
    )

    # Relative reads = proportion of each order within its group
    aggregated_read_data["relative_reads"] = (
        aggregated_read_data["read_count"] / aggregated_read_data["read_sum"]
    )

    # --- Wilcoxon signed-rank test: aliquot 1 vs. aliquot 2 per primer ---
    # Global test across all orders per primer. Compositional dependence between
    # orders is accepted here; per-order tests showed that the only significant
    # differences stem from stochastic sequencing noise at low-abundance taxa
    # (Hymenoptera), not from a systematic aliquot effect.
    PRIMER_LABELS = {"fwh2": "fwh2", "bf3": "BF3"}
    print("\n=== Metabarcoding aliquot consistency (Wilcoxon signed-rank test) ===")
    for primer, sub in aggregated_read_data.groupby("primer", sort=False):
        aliquot_1 = sub[sub["aliquot"] == 1].set_index(["sample", "cycles", "order"])[
            "relative_reads"
        ]
        aliquot_2 = sub[sub["aliquot"] == 2].set_index(["sample", "cycles", "order"])[
            "relative_reads"
        ]

        paired = pd.DataFrame({"a1": aliquot_1, "a2": aliquot_2}).dropna()
        stat, p = wilcoxon(paired["a1"], paired["a2"])
        label = PRIMER_LABELS.get(primer, primer)
        print(f"  {label}: statistic={stat:.2f}, p={p:.4e}, n_pairs={len(paired)}")

    plot_aliquot_consistency(aggregated_read_data)

    # --- Pool aliquots into a single value per (sample, cycles, primer, order) ---
    # Summing read_count and read_sum across both aliquots gives the pooled
    # proportion (weighted by sequencing depth), not the arithmetic mean.
    aggregated_read_data = (
        aggregated_read_data.groupby(by=["sample", "cycles", "primer", "order"])
        .sum()[["read_count", "read_sum"]]
        .reset_index()
    )

    # Recompute relative reads from pooled counts
    aggregated_read_data["relative_reads"] = (
        aggregated_read_data["read_count"] / aggregated_read_data["read_sum"]
    )

    return aggregated_read_data


def plot_aliquot_consistency(aggregated_read_data: pd.DataFrame) -> None:
    """Plot extraction aliquot reproducibility (Supporting Information 7).

    For each primer, creates a scatter plot comparing aliquot 1 vs. aliquot 2
    relative read proportions for every taxon and sample. Points lying on the
    1:1 diagonal indicate that both aliquots gave the same result. Points far
    from the diagonal indicate extraction variability for that taxon.
    Saved as Supporting Information 7.
    """
    PRIMER_LABELS = {"fwh2": "fwh2", "bf3": "BF3"}

    fig, axes = plt.subplots(1, 2, figsize=(12, 6))

    for ax, (primer, label) in zip(axes, PRIMER_LABELS.items()):
        # Pivot so each row is one (sample, order, cycles) pair with
        # aliquot 1 and aliquot 2 as separate columns
        sub = aggregated_read_data[aggregated_read_data["primer"] == primer]
        pivot = sub.pivot_table(
            index=["sample", "order", "cycles"],
            columns="aliquot",
            values="relative_reads",
        ).reset_index()

        # Show legend only on the last panel to avoid duplicates
        show_legend = primer == "bf3"
        sns.scatterplot(
            data=pivot,
            x=1,
            y=2,
            hue="order",
            palette=ORDER_PALETTE,
            hue_order=ORDER_NAMES,
            edgecolor="black",
            linewidth=1,
            legend=show_legend,
            ax=ax,
        )

        # 1:1 reference line — perfect aliquot agreement
        ax.plot([0, 1], [0, 1], linestyle="--", color="black")
        ax.set_xlim(0, 1)
        ax.set_ylim(0, 1)
        ax.set_aspect("equal")
        ax.set_xlabel("Relative reads in aliquot 1")
        ax.set_ylabel("Relative reads in aliquot 2")
        ax.set_title(label)

    # Move the seaborn legend from the right panel to a shared, centred position
    handles, labels = axes[-1].get_legend_handles_labels()
    axes[-1].get_legend().remove()
    fig.legend(
        handles=handles,
        labels=labels,
        loc="lower center",
        ncol=len(ORDER_NAMES),
        frameon=True,
        handletextpad=0.3,
        bbox_to_anchor=(0.5, 0.02),
    )

    fig.suptitle("Aliquot consistency of relative read proportions")
    fig.subplots_adjust(bottom=0.15)
    plt.savefig("supporting information 7.pdf", dpi=300, bbox_inches="tight")
    plt.close(fig)


def aggregate_ddpcr_data(supporting_information_5: Path) -> pd.DataFrame:
    """Load ddPCR copy number data, verify replicate agreement, and pool replicates.

    Droplet digital PCR (ddPCR) gives an absolute count of mitochondrial DNA
    (mtDNA) copies per taxon in each sample. This function loads those raw
    counts (Supporting Information 5), converts them to relative proportions
    (each taxon's copies divided by total copies in that sample), and checks
    whether two independent ddPCR replicates (A and B) agree using a Wilcoxon
    signed-rank test. A scatter plot comparing the replicates is saved as
    Supporting Information 8. The two replicates are then averaged to obtain
    a single reliable copy-number estimate per sample and taxon.
    Returns the pooled copy-number table.
    """
    # --- Load ddPCR data ---
    ddpcr_data = pd.read_excel(supporting_information_5)

    # --- Compute relative copy numbers per (sample, replicate) ---
    total_copies = (
        ddpcr_data.groupby(by=["sample", "replicate"])
        .sum()["mtDNA copies"]
        .reset_index()
        .rename(columns={"mtDNA copies": "total copies"})
    )

    ddpcr_data = ddpcr_data.merge(
        total_copies,
        how="left",
        on=["sample", "replicate"],
    )

    ddpcr_data["relative copies"] = (
        ddpcr_data["mtDNA copies"] / ddpcr_data["total copies"]
    )

    # --- Scatterplot: replicate A vs. replicate B (supporting information 8) ---
    pivot = ddpcr_data.pivot_table(
        index=["sample", "order"],
        columns="replicate",
        values="relative copies",
    ).reset_index()

    fig, ax = plt.subplots(figsize=(6, 6))
    sns.scatterplot(
        data=pivot,
        x="A",
        y="B",
        hue="order",
        palette=ORDER_PALETTE,
        hue_order=ORDER_NAMES,
        edgecolor="black",
        linewidth=1,
        ax=ax,
    )
    ax.plot([0, 1], [0, 1], linestyle="--", color="black")
    ax.set_xlim(0, 1)
    ax.set_ylim(0, 1)
    ax.set_aspect("equal")
    ax.set_xlabel("Relative mtDNA copies in replicate A")
    ax.set_ylabel("Relative mtDNA copies in replicate B")
    ax.set_title("Replicate consistency of relative mtDNA copy numbers")
    ax.legend(frameon=True, handletextpad=0.3)
    plt.savefig("supporting information 8.pdf", dpi=300, bbox_inches="tight")
    plt.close(fig)

    # --- Wilcoxon signed-rank test: replicate A vs. replicate B ---
    paired = pivot.dropna(subset=["A", "B"])
    stat, p = wilcoxon(paired["A"], paired["B"])
    print(
        f"\n=== ddPCR replicate consistency (Wilcoxon signed-rank test) ===\n"
        f"  statistic={stat:.2f}, p={p:.4e}, n_pairs={len(paired)}"
    )

    # --- Pool replicates by averaging copy numbers per (sample, order) ---
    # Mean (not sum) to preserve the physical scale of copy numbers
    ddpcr_data = (
        ddpcr_data.groupby(by=["sample", "order"])
        .mean(numeric_only=True)[["mtDNA copies"]]
        .reset_index()
    )

    # Recompute totals and relative proportions from pooled means
    total_copies = (
        ddpcr_data.groupby("sample")["mtDNA copies"].sum().rename("total copies")
    )
    ddpcr_data = ddpcr_data.merge(total_copies, on="sample")
    ddpcr_data["relative copies"] = (
        ddpcr_data["mtDNA copies"] / ddpcr_data["total copies"]
    )

    return ddpcr_data


def final_aggregation(
    relative_read_data: pd.DataFrame,
    ddpcr_data: pd.DataFrame,
    supporting_information_1: Path,
) -> pd.DataFrame:
    """Combine the three data sources into one master table.

    This function brings together the three measurements collected for each
    taxon in each sample: relative metabarcoding reads (proportions of DNA
    sequences), relative mtDNA copy numbers from ddPCR, and biomass from
    weighing. After merging, every row in the resulting table holds all three
    values for one taxon in one sample at one PCR cycle count.
    This master table (Supporting Information 9) is the basis for all
    subsequent comparisons and figures.
    """
    supporting_information_1 = pd.read_excel(supporting_information_1)

    # Merge reads ← ddPCR ← biomass on shared (sample, order) keys
    full_results = relative_read_data.merge(
        ddpcr_data, on=["sample", "order"], how="left"
    ).merge(supporting_information_1, on=["sample", "order"], how="left")

    full_results = full_results.sort_values(
        by=["primer", "sample", "cycles", "order"],
        ascending=[False, True, True, True],
    )

    return full_results


def plot_stacked_barplots(
    full_results: pd.DataFrame, primer: str, cycles: int, filename: str
) -> None:
    """Plot per-sample community compositions as stacked bar charts (SI 11/12).

    Creates a 9×9 grid (one panel per sample) where each panel shows three
    stacked bars side by side: relative biomass, relative mtDNA copies, and
    relative metabarcoding reads. Each segment within a bar is coloured by
    insect order. This allows visual comparison of how well the three
    measurement types agree on the relative abundances within each community.
    Saved as Supporting Information 12 (fwh2) and 13 (BF3).
    """
    sub = full_results[
        (full_results["primer"] == primer) & (full_results["cycles"] == cycles)
    ]
    samples = sorted(sub["sample"].unique())

    bar_columns = ["relative biomass", "relative copies", "relative_reads"]
    bar_labels = ["Biomass", "mtDNA copies", "Metabarcoding reads"]
    bar_width = 0.4
    x = np.arange(len(bar_columns)) * bar_width

    fig, axes = plt.subplots(9, 9, figsize=(8, 12), sharey=True)

    for idx, ax in enumerate(axes.flat):
        if idx >= len(samples):
            ax.set_visible(False)
            continue

        sample = samples[idx]
        sample_data = sub[sub["sample"] == sample].set_index("order")
        sample_data = sample_data.reindex(ORDER_NAMES).fillna(0)

        # Stack bars by order, accumulating bottom offsets
        bottom = np.zeros(len(bar_columns))
        for order, color in zip(ORDER_NAMES, ORDER_COLORS):
            values = [sample_data.loc[order, col] for col in bar_columns]
            ax.bar(
                x,
                values,
                bottom=bottom,
                color=color,
                width=bar_width,
                edgecolor="black",
                linewidth=0.3,
            )
            bottom += values

        ax.set_title(f"{sample}", fontsize=7, pad=2)
        ax.set_ylim(0, 1)
        ax.set_xlim(x[0] - bar_width / 2, x[-1] + bar_width / 2)
        ax.tick_params(labelsize=5, length=2)

        # y-axis label only on leftmost column
        if idx % 9 == 0:
            ax.set_ylabel("Proportion", fontsize=6)
        else:
            ax.tick_params(labelleft=False)

        # x-axis labels only on bottom row
        if idx >= 72:
            ax.set_xticks(x)
            ax.set_xticklabels(bar_labels, rotation=45, ha="right", fontsize=5)
        else:
            ax.set_xticks([])

    # Shared legend at bottom
    PRIMER_LABELS = {"fwh2": "fwh2", "bf3": "BF3"}
    primer_label = PRIMER_LABELS.get(primer, primer)
    fig.suptitle(
        f"Per-sample community composition ({primer_label}, {cycles} cycles)",
        fontsize=10,
    )
    handles = [
        plt.Line2D(
            [0],
            [0],
            marker="o",
            color="w",
            markerfacecolor=ORDER_PALETTE[name],
            markeredgecolor="black",
            markeredgewidth=0.3,
            markersize=6,
            label=name,
        )
        for name in ORDER_NAMES
    ]
    fig.legend(
        handles=handles,
        loc="lower center",
        ncol=len(ORDER_NAMES),
        frameon=True,
        handletextpad=0.3,
        bbox_to_anchor=(0.5, -0.03),
        fontsize=7,
    )
    fig.subplots_adjust(hspace=0.4, wspace=0.05, bottom=0.06, top=0.95)
    plt.savefig(filename, dpi=300, bbox_inches="tight")
    plt.close(fig)


def plot_reads_per_cycle(
    full_results: pd.DataFrame, primer: str, filename: str
) -> None:
    """Plot how relative read proportions change across PCR cycle counts (SI 14).

    Creates a 9×9 grid (one panel per sample) where the x-axis shows PCR
    cycle count (6, 8, 10, … 20) and each stacked bar shows the relative
    read proportion of each taxon at that cycle count. If amplification bias
    is cycle-independent (as our constant bias model assumes), the bar
    compositions should look the same at every cycle — any taxon that is
    over- or underrepresented at c=20 should be equally over- or
    underrepresented at c=6. Saved as Supporting Information 14.
    """
    sub = full_results[full_results["primer"] == primer].copy()
    samples = sorted(sub["sample"].unique())
    cycles = sorted(sub["cycles"].unique())

    bar_width = 0.6
    x_pos = np.arange(len(cycles))

    fig, axes = plt.subplots(9, 9, figsize=(14, 14), sharey=True)

    for idx, ax in enumerate(axes.flat):
        if idx >= len(samples):
            ax.set_visible(False)
            continue

        sample = samples[idx]
        sd = sub[sub["sample"] == sample]

        for cx, c in enumerate(cycles):
            cd = sd[sd["cycles"] == c].set_index("order")
            bottom = 0.0
            for order, color in zip(ORDER_NAMES, ORDER_COLORS):
                val = cd.loc[order, "relative_reads"] if order in cd.index else 0.0
                ax.bar(
                    cx,
                    val,
                    bottom=bottom,
                    color=color,
                    width=1.0,
                    edgecolor="black",
                    linewidth=0.3,
                )
                bottom += val

        ax.set_title(f"{sample}", fontsize=6, pad=2)
        ax.set_ylim(0, 1)
        ax.set_xlim(-0.5, len(cycles) - 0.5)
        ax.tick_params(labelsize=4, length=2)

        if idx % 9 == 0:
            ax.set_ylabel("Rel. reads", fontsize=5)
        else:
            ax.tick_params(labelleft=False)

        if idx >= 72:
            ax.set_xticks(x_pos)
            ax.set_xticklabels(
                [str(c) for c in cycles], fontsize=4, rotation=45, ha="right"
            )
        else:
            ax.set_xticks([])

    PRIMER_LABELS = {"fwh2": "fwh2", "bf3": "BF3"}
    primer_label = PRIMER_LABELS.get(primer, primer)
    fig.suptitle(
        f"Relative metabarcoding read proportions across PCR cycles ({primer_label})",
        fontsize=10,
    )
    handles = [
        plt.Line2D(
            [0],
            [0],
            marker="o",
            color="w",
            markerfacecolor=ORDER_PALETTE[name],
            markeredgecolor="black",
            markeredgewidth=0.3,
            markersize=6,
            label=name,
        )
        for name in ORDER_NAMES
    ]
    fig.legend(
        handles=handles,
        loc="lower center",
        ncol=len(ORDER_NAMES),
        frameon=True,
        handletextpad=0.3,
        bbox_to_anchor=(0.5, -0.01),
        fontsize=7,
    )
    fig.subplots_adjust(hspace=0.45, wspace=0.05, bottom=0.05, top=0.95)
    plt.savefig(filename, dpi=300, bbox_inches="tight")
    plt.close(fig)


def compute_fold_changes_and_ranks(full_results: pd.DataFrame) -> pd.DataFrame:
    """Calculate fold-changes and abundance ranks between the three measurement types.

    To quantify how well metabarcoding reads track biomass and mtDNA copies,
    this function computes log2 fold-changes between all three pairs of
    measurements (biomass vs. copies, copies vs. reads, biomass vs. reads).
    A fold-change of 0 means perfect agreement; +1 means the second measure
    is twice as high as the first (2-fold over-representation); -1 means
    half as high (2-fold under-representation). The ±log2(1.5) band
    corresponds to a ±1.5× agreement zone. Additionally, within each sample
    the taxa are ranked by abundance (rank 1 = most abundant) for each
    measurement type, which allows checking whether the rank order is
    preserved even when absolute values differ.
    """

    # --- Log2 fold-changes between all three measurement pairs ---
    def safe_log2_ratio(numerator, denominator):
        ratio = numerator / denominator
        return np.log2(ratio.replace(0, np.nan))

    full_results["fc_biomass_copies"] = safe_log2_ratio(
        full_results["relative copies"], full_results["relative biomass"]
    )
    full_results["fc_copies_reads"] = safe_log2_ratio(
        full_results["relative_reads"], full_results["relative copies"]
    )
    full_results["fc_biomass_reads"] = safe_log2_ratio(
        full_results["relative_reads"], full_results["relative biomass"]
    )

    # --- Within-sample ranks (1 = highest proportion) ---
    group_keys = ["sample", "cycles", "primer"]
    full_results["rank_biomass"] = full_results.groupby(group_keys)[
        "relative biomass"
    ].rank(ascending=False)
    full_results["rank_copies"] = full_results.groupby(group_keys)[
        "relative copies"
    ].rank(ascending=False)
    full_results["rank_reads"] = full_results.groupby(group_keys)[
        "relative_reads"
    ].rank(ascending=False)

    return full_results


def export_table1_rank_concordance(
    full_results: pd.DataFrame, filename: str = "table 1.xlsx"
) -> None:
    """Export Table 1: how often do the three measurement types agree on rank order?

    For each pair of measurements (biomass vs. copies, copies vs. reads,
    biomass vs. reads) and each primer, counts how many taxa have the same
    within-sample rank (e.g. most abundant in both biomass and reads). Reports
    the count and percentage of rank-concordant cases. Saved as Table 1.
    """
    rank_pairs = [
        ("rank_biomass", "rank_copies", "Biomass - Copies"),
        ("rank_copies", "rank_reads", "Copies - Reads"),
        ("rank_biomass", "rank_reads", "Biomass - Reads"),
    ]
    sub = full_results[full_results["cycles"] == 20]
    rows = []
    for primer, primer_label in [("fwh2", "fwh2"), ("bf3", "BF3")]:
        p = sub[sub["primer"] == primer]
        row = {"Primer": primer_label}
        for rank_a, rank_b, label in rank_pairs:
            match = int((p[rank_a] == p[rank_b]).sum())
            total = len(p)
            pct = match / total * 100
            row[label] = f"{match}/{total} ({pct:.1f}%)"
        rows.append(row)
    pd.DataFrame(rows).to_excel(filename, index=False)


def export_si11_over_under(
    full_results: pd.DataFrame, filename: str = "supporting information 11.xlsx"
) -> None:
    """Export Supporting Information 11: over- and under-representation counts per taxon.

    For each pair of measurements and each insect order, counts how many
    samples fall into three categories: over-represented (fold-change > +1.5×),
    concordant (within ±1.5×), or under-represented (fold-change < -1.5×).
    The results are written into three Excel sheets — one sheet per measurement
    pair — with counts and percentages for both primers side by side.
    Note: Biomass vs. copies does not involve sequencing reads and is therefore
    primer-independent; the values are identical for fwh2 and BF3.
    """
    fc_pairs = [
        ("fc_biomass_copies", "Biomass vs Copies"),
        ("fc_copies_reads", "Copies vs Reads"),
        ("fc_biomass_reads", "Biomass vs Reads"),
    ]
    band = np.log2(1.5)
    sub = full_results[full_results["cycles"] == 20]

    with pd.ExcelWriter(filename, engine="openpyxl") as writer:
        for fc_col, sheet_name in fc_pairs:
            # Biomass vs Copies is primer-independent — one table using fwh2 rows.
            # Copies/Reads and Biomass/Reads show fwh2 and BF3 side by side.
            primers = (
                [("fwh2", "fwh2")]
                if fc_col == "fc_biomass_copies"
                else [("fwh2", "fwh2"), ("bf3", "BF3")]
            )
            rows = []
            for order in ORDER_NAMES:
                row = {"Order": order}
                for primer, col_prefix in primers:
                    vals = sub[(sub["primer"] == primer) & (sub["order"] == order)][
                        fc_col
                    ].dropna()
                    n = len(vals)
                    over = int((vals > band).sum())
                    conc = int((vals.abs() <= band).sum())
                    under = int((vals < -band).sum())
                    prefix = "" if fc_col == "fc_biomass_copies" else f"{col_prefix} "
                    row[f"{prefix}n"] = n
                    row[f"{prefix}over (n)"] = over
                    row[f"{prefix}over (%)"] = round(over / n * 100, 1) if n else 0
                    row[f"{prefix}conc. (n)"] = conc
                    row[f"{prefix}conc. (%)"] = round(conc / n * 100, 1) if n else 0
                    row[f"{prefix}under (n)"] = under
                    row[f"{prefix}under (%)"] = round(under / n * 100, 1) if n else 0
                rows.append(row)
            pd.DataFrame(rows).to_excel(writer, sheet_name=sheet_name, index=False)


def plot_fold_change_stripplot(
    full_results: pd.DataFrame, primer: str, cycles: int, filename: str
) -> None:
    """Plot fold-changes between measurement types per sample (Figure 2).

    Creates Figure 2 with three panels, one for each pair of measurements
    (A: biomass vs. mtDNA copies, B: copies vs. reads, C: biomass vs. reads).
    Each dot represents one taxon in one sample, coloured by insect order.
    The grey band marks the ±1.5× agreement zone — dots inside the band are
    considered concordant. Dots above the band indicate the second measurement
    is more than 1.5× higher; dots below indicate more than 1.5× lower.
    Saved as Figure 2.
    """
    sub = full_results[
        (full_results["primer"] == primer) & (full_results["cycles"] == cycles)
    ].copy()
    sub["sample"] = sub["sample"].astype(str)
    sample_order = [str(s) for s in sorted(full_results["sample"].unique())]

    panels = [
        ("fc_biomass_copies", "A)  Biomass vs. mtDNA copies"),
        ("fc_copies_reads", "B)  mtDNA copies vs. metabarcoding reads"),
        ("fc_biomass_reads", "C)  Biomass vs. metabarcoding reads"),
    ]

    # ±1.5x band (x1.5 / ÷1.5) symmetric in log2 space
    band = np.log2(1.5)

    fig, axes = plt.subplots(3, 1, figsize=(16, 10), sharex=True)

    for ax, (fc_col, title) in zip(axes, panels):
        # Agreement band and reference lines
        ax.axhspan(-band, band, color="lightgray", alpha=0.4, zorder=0)
        ax.axhline(0, color="gray", linestyle="-", linewidth=0.8, zorder=1)
        ax.axhline(band, color="silver", linestyle="--", linewidth=0.6, zorder=1)
        ax.axhline(-band, color="silver", linestyle="--", linewidth=0.6, zorder=1)

        sns.stripplot(
            data=sub,
            x="sample",
            y=fc_col,
            hue="order",
            palette=ORDER_PALETTE,
            hue_order=ORDER_NAMES,
            order=sample_order,
            dodge=False,
            size=6,
            alpha=0.8,
            edgecolor="black",
            linewidth=0.3,
            legend=False,
            ax=ax,
            zorder=2,
        )

        ax.set_title(title, fontsize=14)
        ax.set_ylabel("Fold-change (log$_2$)", fontsize=12)
        ax.set_xlabel("")
        ax.tick_params(labelsize=10)

    # x-axis labels only on bottom panel
    axes[-1].set_xticklabels(sample_order, rotation=90, fontsize=10)
    axes[-1].set_xlabel("Sample", fontsize=12)

    # Shared legend at bottom (scatter markers, matching SI 7 style)
    handles = [
        plt.Line2D(
            [0],
            [0],
            marker="o",
            color="w",
            markerfacecolor=ORDER_PALETTE[name],
            markeredgecolor="black",
            markeredgewidth=0.3,
            markersize=6,
            label=name,
        )
        for name in ORDER_NAMES
    ]
    fig.legend(
        handles=handles,
        loc="lower center",
        ncol=len(ORDER_NAMES),
        frameon=True,
        handletextpad=0.3,
        bbox_to_anchor=(0.5, 0.0),
        fontsize=11,
    )

    fig.subplots_adjust(hspace=0.18, bottom=0.10)
    # Save as both PDF and PNG
    base = filename.rsplit(".", 1)[0]
    plt.savefig(f"{base}.pdf", dpi=300, bbox_inches="tight")
    plt.savefig(f"{base}.png", dpi=300, bbox_inches="tight")
    plt.close(fig)


def compute_cycle_calibration(
    full_results: pd.DataFrame, primer: str = "fwh2"
) -> pd.DataFrame:
    """Estimate amplification bias using the cycle calibration approach of Shelton et al. (2023).

    The key idea: if amplification efficiency differs between taxa, the
    log-ratio of their read counts changes linearly with PCR cycle number.
    By measuring the same sample at many different cycle counts (here 8 counts:
    c = 6, 8, 10, … 20), we can fit a straight line to log(reads_i / reads_ref)
    vs. cycle number for each taxon i. The slope of this line estimates the
    efficiency difference (E_cycle_cal), and the y-intercept estimates the
    log-ratio of starting copy numbers before PCR — which is what we actually
    want to recover. Converting those intercepts back to proportions gives
    the predicted relative copy numbers (pred_copies_cycle_cal).

    Crucially, this approach requires no external calibration samples: all
    information is extracted from the variation across cycle counts within
    each community sample. The most abundant taxon at c=20 is used as the
    reference (choice of reference does not affect the predicted proportions).
    Results are returned as a table with one row per sample and taxon,
    later exported as Supporting Information 15.
    """
    sub = full_results[full_results["primer"] == primer].copy()

    records = []
    for sample in sorted(sub["sample"].unique()):
        sample_data = sub[sub["sample"] == sample]
        orders_in_sample = sample_data["order"].unique()
        all_cycles = sorted(sample_data["cycles"].unique())

        if len(orders_in_sample) < 2 or len(all_cycles) < 3:
            continue

        # Reference taxon: most abundant at c=20 (choice is arbitrary,
        # predicted proportions are identical regardless of reference)
        c20_data = sample_data[sample_data["cycles"] == 20]
        if c20_data.empty:
            continue
        ref_order = c20_data.loc[c20_data["read_count"].idxmax(), "order"]

        # Fit log-ratio regression for each non-reference taxon
        betas = {ref_order: 0.0}
        alphas = {ref_order: 0.0}
        for order in orders_in_sample:
            if order == ref_order:
                continue
            log_ratios = []
            valid_cycles = []
            for c in all_cycles:
                c_data = sample_data[sample_data["cycles"] == c]
                ri = c_data.loc[c_data["order"] == order, "read_count"].values
                rr = c_data.loc[c_data["order"] == ref_order, "read_count"].values
                if len(ri) and len(rr) and ri[0] > 0 and rr[0] > 0:
                    log_ratios.append(np.log(ri[0] / rr[0]))
                    valid_cycles.append(c)
            if len(valid_cycles) >= 3:
                result = linregress(valid_cycles, log_ratios)
                betas[order] = result.intercept
                alphas[order] = result.slope
            else:
                betas[order] = np.nan
                alphas[order] = np.nan

        # Convert intercepts to predicted proportions via softmax
        valid_orders = [
            o for o in orders_in_sample if not np.isnan(betas.get(o, np.nan))
        ]
        if len(valid_orders) < 2:
            continue

        exp_betas = {o: np.exp(betas[o]) for o in valid_orders}
        total_eb = sum(exp_betas.values())

        for o in valid_orders:
            records.append(
                {
                    "sample": sample,
                    "order": o,
                    "ref_species": ref_order,
                    "E_cycle_cal": np.exp(alphas[o]),
                    "pred_copies_cycle_cal": exp_betas[o] / total_eb,
                }
            )

    return pd.DataFrame(records)


def predict_copies_model_comparison(full_results: pd.DataFrame) -> None:
    """Compare four approaches to estimating taxon abundances from metabarcoding reads.

    The central question of this study: can we correct for amplification bias
    in metabarcoding data to obtain better estimates of true taxon abundances
    (measured here as relative mtDNA copy numbers by ddPCR)?

    Four approaches are compared:
      1. No correction — use raw read proportions as the abundance estimate.
         This is the standard metabarcoding output, no additional data needed.
      2. Cycle calibration (Shelton et al. 2023) — uses the change in read
         ratios across PCR cycle counts within each sample to infer starting
         copy proportions. No external calibration data needed.
      3. Cycle-dependent bias model (Shelton exponential model) — uses 14
         two-species calibration communities to estimate a per-cycle efficiency
         factor E_i for each taxon. Correction is applied as reads / E_i^c,
         so the correction strength changes with cycle count.
      4. Constant bias model (this study) — also uses the 14 calibration
         communities, but assumes the bias is the same at every cycle count.
         Correction is applied as reads / E_i (no cycle dependency).

    The 14 calibration communities each contain exactly one target taxon plus
    Blattodea as the reference (Blattodea E_i = 1 by definition). The
    remaining 67 communities are used as out-of-sample test data for approaches
    3 and 4. Both E_i values are estimated from measurements at c=20 only,
    making the comparison fair.

    Outputs: Figure 4, Table 2, Supporting Information 18, 19, and 20.
    """
    primer = "fwh2"
    sub = full_results[full_results["primer"] == primer].copy()

    # ----------------------------------------------------------------
    # Step 1: Identify calibration communities and estimate E_i
    # ----------------------------------------------------------------
    # The 14 calibration communities each contain exactly two taxa:
    # one target insect order plus Blattodea as the reference.
    estimation_samples = []
    for sample, grp in sub[sub["cycles"] == 20].groupby("sample"):
        orders = grp["order"].unique()
        if len(orders) == 2 and "Blattodea" in orders:
            estimation_samples.append(sample)

    # For each calibration sample, compute the odds ratio at c=20:
    #   odds = (reads_i / reads_Blattodea) / (copies_i / copies_Blattodea)
    # This measures how much taxon i is over- or under-represented in
    # reads relative to what its copy numbers would predict.
    # Constant bias model: E_i = mean(odds) — the same factor at every cycle.
    # Cycle-dependent model: E_i = mean(odds^(1/20)) — a per-cycle factor
    #   that, when raised to the power of cycle count c, gives the total bias.
    # Both are calibrated from the same data (c=20 only) for a fair comparison.
    e_orig_per_taxon = {}
    e_exp_per_taxon = {}
    for sample in estimation_samples:
        sd = sub[(sub["sample"] == sample) & (sub["cycles"] == 20)].set_index("order")
        other = [o for o in sd.index if o != "Blattodea"][0]
        rr_i = sd.loc[other, "relative_reads"]
        rc_i = sd.loc[other, "relative copies"]
        rr_b = sd.loc["Blattodea", "relative_reads"]
        rc_b = sd.loc["Blattodea", "relative copies"]
        if rr_i == 0 or rr_i == 1 or rc_i == 0:
            continue
        odds = (rr_i * rc_b) / (rc_i * rr_b)
        e_orig_per_taxon.setdefault(other, []).append(odds)
        e_exp_per_taxon.setdefault(other, []).append(odds ** (1.0 / 20))

    e_orig = {o: np.mean(v) for o, v in e_orig_per_taxon.items()}
    e_orig["Blattodea"] = 1.0  # Blattodea is the reference; its E_i is 1 by definition
    e_exp = {o: np.mean(v) for o, v in e_exp_per_taxon.items()}
    e_exp["Blattodea"] = 1.0

    # ----------------------------------------------------------------
    # Step 1b: Estimate E_i for BF3 (constant bias model, c=20 only)
    # ----------------------------------------------------------------
    # Uses the same 14 calibration communities but the BF3 read columns.
    sub_bf3 = full_results[full_results["primer"] == "bf3"].copy()
    e_orig_bf3_per_taxon = {}
    for sample in estimation_samples:
        sd = sub_bf3[
            (sub_bf3["sample"] == sample) & (sub_bf3["cycles"] == 20)
        ].set_index("order")
        if len(sd) < 2 or "Blattodea" not in sd.index:
            continue
        other = [o for o in sd.index if o != "Blattodea"][0]
        rr_i = sd.loc[other, "relative_reads"]
        rc_i = sd.loc[other, "relative copies"]
        rr_b = sd.loc["Blattodea", "relative_reads"]
        rc_b = sd.loc["Blattodea", "relative copies"]
        if rr_i == 0 or rr_i == 1 or rc_i == 0:
            continue
        odds = (rr_i * rc_b) / (rc_i * rr_b)
        e_orig_bf3_per_taxon.setdefault(other, []).append(odds)
    e_orig_bf3 = {o: np.mean(v) for o, v in e_orig_bf3_per_taxon.items()}
    e_orig_bf3["Blattodea"] = 1.0

    # ----------------------------------------------------------------
    # Step 2a: Run cycle calibration (Shelton et al. 2023) for all samples
    # ----------------------------------------------------------------
    # This approach does not need external calibration communities —
    # it uses within-sample variation across cycle counts instead.
    cycle_cal_table = compute_cycle_calibration(full_results, primer=primer)
    cycle_cal_n0 = {
        (row.sample, row.order): row.pred_copies_cycle_cal
        for row in cycle_cal_table.itertuples()
    }

    # ----------------------------------------------------------------
    # Step 2b: Compute predictions for all four approaches
    # ----------------------------------------------------------------
    # prediction_samples: the 67 communities not used for calibration.
    # Bias-corrected models (3 & 4) are only evaluated on these samples
    # to ensure the comparison is out-of-sample (i.e. the model has never
    # seen these communities during calibration).
    prediction_samples = set(sub["sample"].unique()) - set(estimation_samples)
    all_samples = set(sub["sample"].unique())

    records = []
    for sample in sorted(all_samples):
        for cycles in sorted(sub["cycles"].unique()):
            sample_data = sub[(sub["sample"] == sample) & (sub["cycles"] == cycles)]
            if sample_data.empty:
                continue

            orders = sample_data["order"].values
            reads = sample_data.set_index("order")["relative_reads"]
            obs_copies = sample_data.set_index("order")["relative copies"]

            for order in orders:
                obs_val = obs_copies[order]

                # Approach 1: no correction — reads used as-is
                records.append(
                    {
                        "sample": sample,
                        "cycles": cycles,
                        "order": order,
                        "method": "no correction",
                        "obs": obs_val,
                        "pred": reads[order],
                        "abs_error": abs(reads[order] - obs_val),
                    }
                )

                # Approach 2: cycle calibration — one predicted proportion per
                # (sample, order), independent of cycle count
                cc_pred = cycle_cal_n0.get((sample, order), np.nan)
                if not np.isnan(cc_pred):
                    records.append(
                        {
                            "sample": sample,
                            "cycles": cycles,
                            "order": order,
                            "method": "cycle calibration",
                            "obs": obs_val,
                            "pred": cc_pred,
                            "abs_error": abs(cc_pred - obs_val),
                        }
                    )

                # Approaches 3 & 4: bias-corrected models, out-of-sample only
                if sample in prediction_samples:
                    # Approach 3 (constant bias): divide each taxon's reads by
                    # its E_i, then re-normalise so proportions sum to 1
                    corr_ne = {o: reads[o] / e_orig.get(o, 1.0) for o in orders}
                    t_ne = sum(corr_ne.values())
                    if t_ne > 0:
                        pred_ne = corr_ne[order] / t_ne
                        records.append(
                            {
                                "sample": sample,
                                "cycles": cycles,
                                "order": order,
                                "method": "non-exponential",
                                "obs": obs_val,
                                "pred": pred_ne,
                                "abs_error": abs(pred_ne - obs_val),
                            }
                        )
                    # Approach 4 (cycle-dependent bias): divide by E_i raised
                    # to the power of the actual cycle count used
                    corr_exp = {
                        o: reads[o] / (e_exp.get(o, 1.0) ** cycles) for o in orders
                    }
                    t_exp = sum(corr_exp.values())
                    if t_exp > 0:
                        pred_exp = corr_exp[order] / t_exp
                        records.append(
                            {
                                "sample": sample,
                                "cycles": cycles,
                                "order": order,
                                "method": "cycle-dependent",
                                "obs": obs_val,
                                "pred": pred_exp,
                                "abs_error": abs(pred_exp - obs_val),
                            }
                        )

    df = pd.DataFrame(records)
    df_common = df[df["sample"].isin(prediction_samples)]

    # ----------------------------------------------------------------
    # Step 5b: Figure 3 — no correction vs. cycle calibration only
    # ----------------------------------------------------------------
    # Both methods require no external calibration data, so all 81
    # communities are used here (not just the 67 out-of-sample ones
    # needed for the fair 3-way comparison above).
    # Shared y-axis since both panels are on the same scale (0-1).
    # RMSE and n are reported in the figure caption, not on the plot.
    figure3_titles = {
        "no correction": "A)  No correction (all cycles)",
        "cycle calibration": "B)  Cycle calibration",
    }

    fig3, axes3 = plt.subplots(1, 2, figsize=(12, 6), sharey=True)
    figure3_methods = ["no correction", "cycle calibration"]

    for ax, method_name in zip(axes3, figure3_methods):
        md = df[df["method"] == method_name]
        if method_name == "cycle calibration":
            md = md.drop_duplicates(subset=["sample", "order"])
        for order in ORDER_NAMES:
            od = md[md["order"] == order]
            if od.empty:
                continue
            ax.scatter(
                od["obs"],
                od["pred"],
                color=ORDER_PALETTE[order],
                label=order,
                alpha=0.5,
                s=15,
                edgecolor="none",
            )
        ax.set_xlim(0, 1)
        ax.set_ylim(0, 1)
        ax.set_aspect("equal")
        ax.set_xlabel("Observed relative copies (ddPCR)")
        ax.set_title(figure3_titles[method_name], fontsize=13)

        obs_v, pred_v = md["obs"].values, md["pred"].values
        ss_res = np.sum((obs_v - pred_v) ** 2)
        ss_tot = np.sum((obs_v - obs_v.mean()) ** 2)
        r2 = 1 - ss_res / ss_tot
        ax.annotate(
            f"R² = {r2:.3f}",
            xy=(0.05, 0.90),
            xycoords="axes fraction",
            fontsize=12,
            fontweight="bold",
            bbox=dict(boxstyle="round", facecolor="white", alpha=0.85),
        )

    axes3[0].set_ylabel("Predicted relative copies")
    axes3[1].legend(fontsize=8, loc="lower right", markerscale=1.5)

    fig3.tight_layout()
    plt.savefig("figure 3.pdf", dpi=300, bbox_inches="tight")
    plt.savefig("figure 3.png", dpi=300, bbox_inches="tight")
    plt.close(fig3)

    # ----------------------------------------------------------------
    # Step 6: Aitchison distance per sample
    # ----------------------------------------------------------------
    # Aitchison distance is the appropriate metric for compositional data
    # (Aitchison 1986). It measures the Euclidean distance between two
    # compositions in centered log-ratio (CLR) space. Unlike R² or RMSE,
    # it respects the multiplicative structure of proportions: a shift
    # from 0.01 to 0.02 (2x) weighs the same as 0.5 to 1.0 (2x).
    # Shelton et al. (2023) use this metric for their model comparison.
    def aitchison_distance(x, y):
        x = np.maximum(np.array(x, dtype=float), 1e-10)
        y = np.maximum(np.array(y, dtype=float), 1e-10)
        clr_x = np.log(x) - np.log(x).mean()
        clr_y = np.log(y) - np.log(y).mean()
        return np.sqrt(np.sum((clr_x - clr_y) ** 2))

    # ----------------------------------------------------------------
    # Aitchison distance across all cycles (4 series)
    # ----------------------------------------------------------------
    aitchison_all_cycles = []
    for sample in sorted(prediction_samples):
        sd20 = sub[(sub["sample"] == sample) & (sub["cycles"] == 20)]
        if sd20.empty:
            continue
        orders = sorted(sd20["order"].values)
        obs_vec = [
            sd20.loc[sd20["order"] == o, "relative copies"].values[0] for o in orders
        ]
        if any(v == 0 for v in obs_vec):
            continue

        cc_vec = [cycle_cal_n0.get((sample, o), np.nan) for o in orders]
        d_cc = (
            np.nan
            if any(np.isnan(v) for v in cc_vec)
            else aitchison_distance(obs_vec, cc_vec)
        )

        for cycles in sorted(sub["cycles"].unique()):
            sd = sub[(sub["sample"] == sample) & (sub["cycles"] == cycles)]
            if sd.empty or set(sd["order"]) != set(orders):
                continue

            nc_vec = [
                sd.loc[sd["order"] == o, "relative_reads"].values[0] for o in orders
            ]
            # Skip (sample, cycle) combinations with a zero-read taxon:
            # 0 -> exactly 0 under correction, extreme CLR floor artifacts.
            if any(v == 0 for v in nc_vec):
                continue

            reads = sd.set_index("order")["relative_reads"]

            corr_ne = {o: reads[o] / e_orig.get(o, 1.0) for o in orders}
            t_ne = sum(corr_ne.values())
            if t_ne == 0:
                continue
            ne_vec = [corr_ne[o] / t_ne for o in orders]

            corr_exp = {o: reads[o] / (e_exp.get(o, 1.0) ** cycles) for o in orders}
            t_exp = sum(corr_exp.values())
            if t_exp == 0:
                continue
            exp_vec = [corr_exp[o] / t_exp for o in orders]

            aitchison_all_cycles.append(
                {
                    "sample": sample,
                    "cycles": cycles,
                    "d_nocorr": aitchison_distance(obs_vec, nc_vec),
                    "d_ne": aitchison_distance(obs_vec, ne_vec),
                    "d_exp": aitchison_distance(obs_vec, exp_vec),
                    "d_cc": d_cc,
                }
            )

    df_ait_all = pd.DataFrame(aitchison_all_cycles)

    print("Median Aitchison distances (prediction samples, all cycles):")
    cc_dedup_med = df_ait_all.drop_duplicates(subset=["sample"])
    print(f"  No correction:              {df_ait_all['d_nocorr'].median():.4f}")
    print(f"  Cycle calibration:          {cc_dedup_med['d_cc'].median():.4f}")
    print(f"  Cycle-dependent bias model: {df_ait_all['d_exp'].median():.4f}")
    print(f"  Constant bias model:        {df_ait_all['d_ne'].median():.4f}")

    sample_order_x = sorted(df_ait_all["sample"].unique())
    x_pos = {s: i for i, s in enumerate(sample_order_x)}

    # ----------------------------------------------------------------
    # Figure 4: 2×2 gridspec + full-width bottom panel
    # Top row: cycle-dependent bias model | constant bias model (scatter)
    # Bottom: 4-series Aitchison distance per sample (all cycles)
    # ----------------------------------------------------------------
    fig4 = plt.figure(figsize=(13, 13))
    gs = fig4.add_gridspec(2, 2, height_ratios=[1, 1], hspace=0.18, wspace=0.06)

    figure4_specs = [
        ("cycle-dependent", "A) Cycle-dependent bias model", gs[0, 0]),
        ("non-exponential", "B) Constant bias model", gs[0, 1]),
    ]
    axes4_top = []
    for method_name, title, gs_pos in figure4_specs:
        ax = fig4.add_subplot(gs_pos)
        axes4_top.append(ax)
        md = df_common[df_common["method"] == method_name]
        for order in ORDER_NAMES:
            od = md[md["order"] == order]
            if od.empty:
                continue
            ax.scatter(
                od["obs"],
                od["pred"],
                color=ORDER_PALETTE[order],
                label=order,
                alpha=0.5,
                s=15,
                edgecolor="none",
            )
        ax.set_xlim(0, 1)
        ax.set_ylim(0, 1)
        ax.set_aspect("equal")
        ax.set_xlabel("Observed relative copies (ddPCR)")
        ax.set_title(title, fontsize=13)

        obs_v, pred_v = md["obs"].values, md["pred"].values
        ss_res = np.sum((obs_v - pred_v) ** 2)
        ss_tot = np.sum((obs_v - obs_v.mean()) ** 2)
        r2 = 1 - ss_res / ss_tot
        ax.annotate(
            f"R² = {r2:.3f}",
            xy=(0.05, 0.90),
            xycoords="axes fraction",
            fontsize=12,
            fontweight="bold",
            bbox=dict(boxstyle="round", facecolor="white", alpha=0.85),
        )

    axes4_top[0].set_ylabel("Predicted relative copies")
    axes4_top[1].legend(fontsize=8, loc="lower right", markerscale=1.5)

    # Bottom panel: Aitchison distance, 4 series
    ax_bottom = fig4.add_subplot(gs[1, :])

    ax_bottom.scatter(
        [x_pos[s] for s in df_ait_all["sample"]],
        df_ait_all["d_nocorr"],
        color="#AAAAAA",
        s=18,
        alpha=0.5,
        edgecolors="none",
        zorder=2,
        label="No correction",
    )
    cc_dedup = df_ait_all.drop_duplicates(subset=["sample"])
    ax_bottom.scatter(
        [x_pos[s] for s in cc_dedup["sample"]],
        cc_dedup["d_cc"],
        color="#E69F00",
        s=30,
        alpha=0.9,
        edgecolors="black",
        linewidths=0.5,
        zorder=4,
        label="Cycle calibration",
    )
    ax_bottom.scatter(
        [x_pos[s] for s in df_ait_all["sample"]],
        df_ait_all["d_exp"],
        color="#009E73",
        s=18,
        alpha=0.5,
        edgecolors="none",
        zorder=3,
        label="Cycle-dependent bias model",
    )
    ax_bottom.scatter(
        [x_pos[s] for s in df_ait_all["sample"]],
        df_ait_all["d_ne"],
        color="#0072B2",
        s=18,
        alpha=0.5,
        edgecolors="none",
        zorder=3,
        label="Constant bias model",
    )

    # Horizontal median lines per method
    for color, col in [
        ("#AAAAAA", "d_nocorr"),
        ("#009E73", "d_exp"),
        ("#0072B2", "d_ne"),
    ]:
        ax_bottom.axhline(
            df_ait_all[col].median(),
            color=color,
            lw=1.5,
            linestyle="--",
            alpha=0.85,
            zorder=1,
        )
    ax_bottom.axhline(
        cc_dedup["d_cc"].median(),
        color="#E69F00",
        lw=1.5,
        linestyle="--",
        alpha=0.85,
        zorder=1,
    )

    ax_bottom.text(
        -0.02,
        1.02,
        "C)",
        transform=ax_bottom.transAxes,
        fontsize=14,
        fontweight="bold",
        va="bottom",
        ha="left",
    )
    ax_bottom.set_ylabel("Aitchison distance to observed copies")
    ax_bottom.set_xticks(range(len(sample_order_x)))
    ax_bottom.set_xticklabels(
        [str(s) for s in sample_order_x], rotation=90, fontsize=10
    )
    ax_bottom.set_xlim(-1, len(sample_order_x))
    ax_bottom.set_ylim(0, None)
    ax_bottom.legend(loc="upper right", fontsize=9)

    plt.savefig("figure 4.pdf", dpi=300, bbox_inches="tight")
    plt.savefig("figure 4.png", dpi=300, bbox_inches="tight")
    plt.close(fig4)

    # ----------------------------------------------------------------
    # Table 2: Mean E_i per taxon for both bias models
    # ----------------------------------------------------------------
    pd.DataFrame(
        [
            {
                "Taxon": order,
                "E_i, cycle-dependent bias": e_exp.get(order, np.nan),
                "E_i, constant bias": e_orig.get(order, np.nan),
            }
            for order in ORDER_NAMES
        ]
    ).to_excel("table 2.xlsx", index=False)

    # ----------------------------------------------------------------
    # Supporting Information 18: scatter per cycle count, 2 models
    # Rows = cycle count (c=6..20), Cols = cycle-dependent | constant bias
    # Prediction samples only (same as Figure 4)
    # ----------------------------------------------------------------
    cycles_list = sorted(sub["cycles"].unique())
    n_cyc = len(cycles_list)
    fig15, axes15 = plt.subplots(
        n_cyc,
        2,
        figsize=(10, n_cyc * 4),
        sharex=True,
        sharey=True,
        gridspec_kw={"hspace": 0.15, "wspace": 0.06},
    )
    si15_col_specs = [
        ("cycle-dependent", "Cycle-dependent bias model"),
        ("non-exponential", "Constant bias model"),
    ]
    for row_idx, c in enumerate(cycles_list):
        for col_idx, (method_name, col_title) in enumerate(si15_col_specs):
            ax = axes15[row_idx, col_idx]
            md = df_common[
                (df_common["method"] == method_name) & (df_common["cycles"] == c)
            ]
            for order in ORDER_NAMES:
                od = md[md["order"] == order]
                if od.empty:
                    continue
                ax.scatter(
                    od["obs"],
                    od["pred"],
                    color=ORDER_PALETTE[order],
                    label=order if (row_idx == 0 and col_idx == 1) else None,
                    alpha=0.5,
                    s=12,
                    edgecolor="none",
                )
            ax.set_xlim(0, 1)
            ax.set_ylim(0, 1)
            ax.set_aspect("equal")
            ax.tick_params(labelsize=7)
            if row_idx == 0:
                ax.set_title(col_title, fontsize=10)
            if row_idx == n_cyc - 1:
                ax.set_xlabel("Observed relative copies (ddPCR)", fontsize=8)
            if col_idx == 0:
                ax.set_ylabel("Predicted", fontsize=8)
                ax.annotate(
                    f"c = {c}",
                    xy=(-0.35, 0.5),
                    xycoords="axes fraction",
                    fontsize=9,
                    fontweight="bold",
                    ha="center",
                    va="center",
                    rotation=90,
                    annotation_clip=False,
                )
            if len(md) > 0:
                obs_v, pred_v = md["obs"].values, md["pred"].values
                ss_res = np.sum((obs_v - pred_v) ** 2)
                ss_tot = np.sum((obs_v - obs_v.mean()) ** 2)
                r2 = 1 - ss_res / ss_tot if ss_tot > 0 else np.nan
                ax.annotate(
                    f"$R^2 = {r2:.2f}$",
                    xy=(0.05, 0.90),
                    xycoords="axes fraction",
                    fontsize=8,
                    bbox=dict(boxstyle="round", facecolor="white", alpha=0.8),
                )
    axes15[0, 1].legend(fontsize=7, loc="lower right", markerscale=1.5)
    plt.savefig("supporting information 18.pdf", dpi=300, bbox_inches="tight")
    plt.savefig("supporting information 18.png", dpi=150, bbox_inches="tight")
    plt.close(fig15)

    # ----------------------------------------------------------------
    # Supporting Information 19: SI 13 extended with bias model predictions
    # Adds 4 columns: E_i per taxon + predicted copies for both models
    # Predictions computed for ALL fwh2 samples (in- and out-of-sample)
    # ----------------------------------------------------------------
    # Bias predictions for all (sample, cycles) in fwh2 data
    pred_exp_all = {}
    pred_ne_all = {}
    for (smp, cyc), grp in sub.groupby(["sample", "cycles"]):
        idx = grp.set_index("order")
        orders_grp = idx.index.tolist()
        corr_exp = {
            o: idx.loc[o, "relative_reads"] / (e_exp.get(o, 1.0) ** cyc)
            for o in orders_grp
        }
        t_exp = sum(corr_exp.values())
        corr_ne = {
            o: idx.loc[o, "relative_reads"] / e_orig.get(o, 1.0) for o in orders_grp
        }
        t_ne = sum(corr_ne.values())
        for o in orders_grp:
            pred_exp_all[(smp, cyc, o)] = corr_exp[o] / t_exp if t_exp > 0 else np.nan
            pred_ne_all[(smp, cyc, o)] = corr_ne[o] / t_ne if t_ne > 0 else np.nan

    ct = cycle_cal_table.copy()
    ct["primer"] = primer
    si16 = full_results.merge(ct, on=["sample", "order", "primer"], how="left")

    # E_i values are fwh2-specific; bf3 rows get NaN
    si16["E_i, cycle-dependent bias"] = np.where(
        si16["primer"] == primer,
        si16["order"].map(e_exp),
        np.nan,
    )
    si16["E_i, constant bias"] = np.where(
        si16["primer"] == primer,
        si16["order"].map(e_orig),
        np.nan,
    )
    si16["pred_copies_cycle_dep_bias"] = [
        pred_exp_all.get((r.sample, r.cycles, r.order), np.nan)
        for r in si16.itertuples()
    ]
    si16["pred_copies_constant_bias"] = [
        pred_ne_all.get((r.sample, r.cycles, r.order), np.nan)
        for r in si16.itertuples()
    ]

    # Aitchison distances: one value per (sample, cycles), repeated across orders
    # Only available for prediction samples (calibration samples get NaN)
    ait_cols = df_ait_all[["sample", "cycles", "d_nocorr", "d_exp", "d_ne", "d_cc"]].rename(
        columns={
            "d_nocorr": "aitchison_no_correction",
            "d_exp":    "aitchison_cycle_dep_bias",
            "d_ne":     "aitchison_constant_bias",
            "d_cc":     "aitchison_cycle_calibration",
        }
    )
    si16 = si16.merge(ait_cols, on=["sample", "cycles"], how="left")

    si16.to_excel("supporting information 19.xlsx", index=False)

    # ----------------------------------------------------------------
    # Supporting Information 20: scatter per insect order, 2 models
    # Rows = order, Cols = cycle-dependent | constant bias
    # Points coloured by cycle count; prediction samples only
    # ----------------------------------------------------------------
    cycles_list_20 = sorted(sub["cycles"].unique())
    cycle_colors = {
        c: plt.cm.viridis(i / max(len(cycles_list_20) - 1, 1))
        for i, c in enumerate(cycles_list_20)
    }

    n_ord = len(ORDER_NAMES)
    fig20, axes20 = plt.subplots(
        n_ord,
        2,
        figsize=(10, n_ord * 4),
        sharex=True,
        sharey=True,
        gridspec_kw={"hspace": 0.15, "wspace": 0.06},
    )
    si20_col_specs = [
        ("cycle-dependent", "Cycle-dependent bias model"),
        ("non-exponential", "Constant bias model"),
    ]
    for row_idx, order in enumerate(ORDER_NAMES):
        for col_idx, (method_name, col_title) in enumerate(si20_col_specs):
            ax = axes20[row_idx, col_idx]
            md = df_common[
                (df_common["method"] == method_name) & (df_common["order"] == order)
            ]
            for c in cycles_list_20:
                cd = md[md["cycles"] == c]
                if cd.empty:
                    continue
                ax.scatter(
                    cd["obs"],
                    cd["pred"],
                    color=cycle_colors[c],
                    label=f"c = {c}" if (row_idx == 0 and col_idx == 1) else None,
                    alpha=0.6,
                    s=12,
                    edgecolor="none",
                )
            ax.set_xlim(0, 1)
            ax.set_ylim(0, 1)
            ax.set_aspect("equal")
            ax.tick_params(labelsize=7)
            if row_idx == 0:
                ax.set_title(col_title, fontsize=10)
            if row_idx == n_ord - 1:
                ax.set_xlabel("Observed relative copies (ddPCR)", fontsize=8)
            if col_idx == 0:
                ax.set_ylabel("Predicted", fontsize=8)
                ax.annotate(
                    order,
                    xy=(-0.35, 0.5),
                    xycoords="axes fraction",
                    fontsize=9,
                    ha="center",
                    va="center",
                    rotation=90,
                    annotation_clip=False,
                )
            if len(md) > 0:
                obs_v, pred_v = md["obs"].values, md["pred"].values
                ss_res = np.sum((obs_v - pred_v) ** 2)
                ss_tot = np.sum((obs_v - obs_v.mean()) ** 2)
                r2 = 1 - ss_res / ss_tot if ss_tot > 0 else np.nan
                ax.annotate(
                    f"$R^2 = {r2:.2f}$",
                    xy=(0.05, 0.90),
                    xycoords="axes fraction",
                    fontsize=8,
                    bbox=dict(boxstyle="round", facecolor="white", alpha=0.8),
                )
    axes20[0, 1].legend(fontsize=7, loc="lower right", markerscale=1.5)
    plt.savefig("supporting information 20.pdf", dpi=300, bbox_inches="tight")
    plt.savefig("supporting information 20.png", dpi=150, bbox_inches="tight")
    plt.close(fig20)

    # ----------------------------------------------------------------
    # Supporting Information 21: E_i comparison table (fwh2 vs BF3)
    # ----------------------------------------------------------------
    si21_rows = [
        {
            "order": order,
            "E_i fwh2 (constant bias model)": round(e_orig.get(order, np.nan), 4),
            "E_i BF3 (constant bias model)": round(e_orig_bf3.get(order, np.nan), 4),
        }
        for order in ORDER_NAMES
    ]
    pd.DataFrame(si21_rows).to_excel("supporting information 21.xlsx", index=False)

    # ----------------------------------------------------------------
    # Supporting Information 22: BF3 scatter — uncorrected vs. corrected
    # ----------------------------------------------------------------
    # Prediction communities for BF3 (same 14 calibration samples excluded)
    sub_bf3 = full_results[full_results["primer"] == "bf3"].copy()
    prediction_samples_bf3 = set(sub_bf3["sample"].unique()) - set(estimation_samples)

    records_bf3 = []
    for sample in sorted(prediction_samples_bf3):
        sample_data = sub_bf3[
            (sub_bf3["sample"] == sample) & (sub_bf3["cycles"] == 20)
        ]
        if sample_data.empty:
            continue
        orders     = sample_data["order"].values
        reads      = sample_data.set_index("order")["relative_reads"]
        obs_copies = sample_data.set_index("order")["relative copies"]
        corr  = {o: reads[o] / e_orig_bf3.get(o, 1.0) for o in orders}
        total = sum(corr.values())
        for order in orders:
            records_bf3.append({
                "sample": sample,
                "order":  order,
                "obs":    obs_copies[order],
                "pred_uncorrected": reads[order],
                "pred_corrected":   corr[order] / total if total > 0 else np.nan,
            })

    df_bf3 = pd.DataFrame(records_bf3)

    def _pred_r2(obs, pred):
        obs, pred = np.array(obs), np.array(pred)
        mask = ~(np.isnan(obs) | np.isnan(pred))
        ss_res = np.sum((obs[mask] - pred[mask]) ** 2)
        ss_tot = np.sum((obs[mask] - obs[mask].mean()) ** 2)
        return 1 - ss_res / ss_tot

    r2_unc_bf3 = _pred_r2(df_bf3["obs"], df_bf3["pred_uncorrected"])
    r2_cor_bf3 = _pred_r2(df_bf3["obs"], df_bf3["pred_corrected"])

    fig22, axes22 = plt.subplots(1, 2, figsize=(10, 5), sharex=True, sharey=True)
    panel_specs22 = [
        ("pred_uncorrected", "No correction",        r2_unc_bf3),
        ("pred_corrected",   "Constant bias model",  r2_cor_bf3),
    ]
    for ax, (col, title, r2_val) in zip(axes22, panel_specs22):
        for order in ORDER_NAMES:
            sub = df_bf3[df_bf3["order"] == order]
            ax.scatter(
                sub["obs"], sub[col],
                color=ORDER_PALETTE[order], label=order,
                s=22, alpha=0.6, linewidths=0,
            )
        lim_max = max(
            df_bf3["obs"].max(),
            df_bf3["pred_corrected"].max(),
            df_bf3["pred_uncorrected"].max(),
        ) * 1.05
        ax.plot([0, lim_max], [0, lim_max], "k--", lw=0.8, zorder=0)
        ax.set_xlim(0, lim_max)
        ax.set_ylim(0, lim_max)
        ax.set_xlabel("Observed relative copies", fontsize=11)
        ax.set_title(title, fontsize=11)
        ax.text(
            0.04, 0.96, f"R² = {r2_val:.2f}",
            transform=ax.transAxes, fontsize=10, fontweight="bold",
            va="top", ha="left",
            bbox=dict(boxstyle="round", facecolor="white", alpha=0.8),
        )
    axes22[0].set_ylabel("Predicted relative copies", fontsize=11)
    axes22[1].legend(
        fontsize=8, loc="lower right", markerscale=1.5,
        frameon=True, edgecolor="black", framealpha=0.9,
    )
    fig22.suptitle("BF3 — constant bias correction", fontsize=12)
    plt.tight_layout()
    plt.savefig("supporting information 22.pdf", dpi=300, bbox_inches="tight")
    plt.savefig("supporting information 22.png", dpi=150, bbox_inches="tight")
    plt.close(fig22)


def main():
    """Run the complete analysis from raw data files to all figures and tables.

    This function orchestrates the entire study in seven steps:

    Step 1 — qPCR primer validation (Supporting Information 4):
        Each primer pair was tested against all five insect orders in single-species
        reactions. The results are shown as a heatmap confirming that fwh2 and BF3
        amplify all five orders (Supporting Information 4).

    Step 2 — Metabarcoding data processing (Supporting Information 7, 9, 11, 12, 14):
        Raw sequencing reads are loaded and tallied per sample, PCR cycle count,
        insect order, and primer pair. Reads are converted to proportions (relative
        reads) so that samples with different total read counts are comparable.
        Two aliquots were sequenced per sample as a reproducibility check; their
        proportions are compared with a Wilcoxon signed-rank test. Aliquots are
        then pooled by averaging.

    Step 3 — ddPCR data processing (Supporting Information 8):
        Digital droplet PCR gives absolute copy numbers per insect taxon. Technical
        replicates are checked for consistency (Wilcoxon test) and then averaged
        to obtain one copy number per (sample, order, primer).

    Step 4 — Merge all measurements:
        Metabarcoding reads, ddPCR copy numbers, and known biomass proportions are
        joined into one unified table. This combined table (Supporting Information 9)
        is the basis for all downstream analyses.

    Step 5 — Fold-change and rank analysis (Figure 2, Table 1, Supporting Information 11):
        For each taxon and sample we compute how far reads deviate from copy numbers
        and from biomass (log2 fold-change). We also check whether the rank order
        of taxa by reads agrees with the rank order by copies (rank concordance).
        Figure 2 shows the fold-change distributions; Table 1 summarises rank
        concordance; Supporting Information 11 counts over- and underrepresentation
        by order.

    Step 6 — Model comparison (Figure 4, Table 2, Supporting Information 18, 19, 20, 21, and 22):
        Four approaches to predicting taxon proportions from reads are compared:
          1. No correction: use reads directly as a proxy for copies.
          2. Cycle calibration (Shelton et al. 2023): fit a per-cycle decay model
             within each community using reads measured at multiple cycle counts.
          3. Cycle-dependent bias model: estimate a per-cycle amplification
             efficiency factor E_i from 14 two-species calibration communities;
             apply as reads / E_i^c, then renormalise.
          4. Constant bias model (our approach): estimate a single amplification
             efficiency factor E_i (the odds ratio at c=20) from the same 14
             calibration communities; apply as reads / E_i, then renormalise.
        Approaches 3 and 4 are evaluated only on the 67 communities that were
        not used for calibration (out-of-sample test).

    Step 7 — Supporting Information 15:
        The cycle calibration predictions (Supporting Information 15) are exported
        as an extended version of Supporting Information 9. BF3 rows have no cycle
        calibration values because that method requires data at multiple cycle counts,
        and BF3 was only sequenced at c=20.
    """
    # --- Input files ---
    supporting_information_1 = Path(
        "supporting information 1.xlsx"
    )  # community compositions + biomass
    supporting_information_2 = Path(
        "supporting information 2.xlsx"
    )  # ddPCR primer/probe sequences
    supporting_information_3 = Path(
        "supporting information 3.xlsx"
    )  # qPCR validation data
    supporting_information_5 = Path(
        "supporting information 5.xlsx"
    )  # ddPCR raw copy numbers
    supporting_information_6 = Path(
        "supporting information 6.parquet.snappy"
    )  # metabarcoding raw reads

    # --- Step 1: qPCR primer validation heatmap ---
    supporting_information_4(supporting_information_3)

    # --- Step 2: Metabarcoding data processing ---
    aggregated_read_data = aggregate_metabarcoding_data(
        supporting_information_1, supporting_information_6
    )
    relative_read_data = absolute_reads_to_relative_reads(aggregated_read_data)

    # --- Step 3: ddPCR data processing ---
    ddpcr_data = aggregate_ddpcr_data(supporting_information_5)

    # --- Step 4: Merge all data sources ---
    full_results = final_aggregation(
        relative_read_data, ddpcr_data, supporting_information_1
    )

    # --- Step 5: Fold-change, rank concordance, and Figure 2 ---
    full_results = compute_fold_changes_and_ranks(full_results)
    plot_fold_change_stripplot(
        full_results, primer="fwh2", cycles=20, filename="figure 2.pdf"
    )

    # --- Step 6: Model comparison (Figure 4, Table 2, SI 15, SI 16) ---
    predict_copies_model_comparison(full_results)

    # --- Step 7: Export full results and additional figures ---

    # Supporting Information 9: the merged table with all measurements
    full_results.to_excel("supporting information 9.xlsx", index=False)

    # Table 1: rank concordance counts (fwh2 and BF3 side by side)
    export_table1_rank_concordance(full_results, filename="table 1.xlsx")

    # Supporting Information 11: over/concordant/underrepresentation by order
    #   Sheet 1 — Biomass vs Copies  (method-independent; one table)
    #   Sheet 2 — Copies vs Reads    (fwh2 and BF3 side by side)
    #   Sheet 3 — Biomass vs Reads   (fwh2 and BF3 side by side)
    export_si11_over_under(full_results, filename="supporting information 11.xlsx")

    # Supporting Information 12 & 12: stacked barplots of community composition
    # at c=20 for fwh2 (SI 11) and BF3 (SI 12)
    plot_stacked_barplots(
        full_results, primer="fwh2", cycles=20, filename="supporting information 12.pdf"
    )
    plot_stacked_barplots(
        full_results, primer="bf3", cycles=20, filename="supporting information 13.pdf"
    )

    # Supporting Information 14: how read proportions change across cycle counts
    plot_reads_per_cycle(
        full_results, primer="fwh2", filename="supporting information 14.pdf"
    )

    # Supporting Information 15: cycle calibration table (fwh2 only).
    # BF3 rows are kept but their cycle-calibration columns are left blank
    # because cycle calibration requires data at multiple cycle counts,
    # and BF3 was only run at c=20.
    cycle_cal_table = compute_cycle_calibration(full_results, primer="fwh2")
    cycle_cal_table["primer"] = "fwh2"
    cycle_cal_export = full_results.merge(
        cycle_cal_table, on=["sample", "order", "primer"], how="left"
    )
    cycle_cal_export.to_excel("supporting information 15.xlsx", index=False)


if __name__ == "__main__":
    main()
```
