## Supporting information 16 for "Characterizing mitochondrial copy number variation and PCR amplification bias as sources of quantitative constraints in DNA metabarcoding"

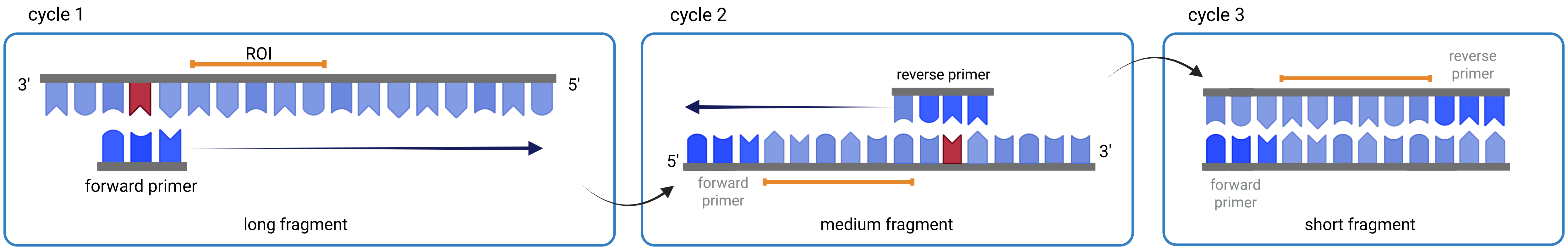


supplementary figure 5: scheme of first three PCR cycles; mismatches (red) are exchanged by matching nucleotides by cycle 3

created with biorender.com
