## Supporting information 17 for "Characterizing mitochondrial copy number variation and PCR amplification bias as sources of quantitative constraints in DNA metabarcoding"

### Supporting information 17: A two-stage model for PCR amplification

#### 1. A two-stage model for PCR amplification

We define a two-stage stochastic model to describe the amplification process of a specific target molecule. Let $N_0$ denote the number of starting molecules. After $c$ PCR cycles, we distinguish two populations of amplicons: $M$ denotes the number of medium-sized fragments amplified with the forward primer only, and $S$ denotes the number of short fragments amplified with both the forward and the reverse primer.
We define $p_1$ as the probability that any starting molecule successfully produces an $M$-fragment in a given cycle, and $p_2$ as the probability that any $M$-fragment successfully produces an $S$-fragment in a given cycle. Both $p_1$ and $p_2$ are assumed to be constant across cycles and independent between molecules. For notational convenience, we write $M(N_{0,c})$ and $S(N_{0,c})$ for the expected total numbers of $M$- and $S$-fragments after $c$ cycles, respectively, making the dependence on $N_0$ explicit in the argument.

#### 2. Expected number of M-fragments

In each cycle, every starting molecule independently produces an $M$-fragment with probability $p_1$. Formally, the number of new $M$-fragments produced in cycle $i$ is a random variable $X_i \sim \mathrm{Bin}(N_0, p_1)$, with $\mathbb{E}[X_i] = N_0 p_1$. Summing the expected per-cycle contributions over all $c$ cycles gives:

$$M(N_{0,c}) = N_0 \cdot c \cdot p_1 \qquad (1)$$

#### 3. Expected number of S-fragments

Once produced, $S$-fragments are no longer subject to primer competition and simply double with each additional PCR cycle. New $S$-fragments are generated from $M$-fragments present at the end of the preceding cycle. This leads to the following recursion:

$$S(N_{0,c}) = p_2 \cdot M(N_{0,c-1}) + 2 \cdot S(N_{0,c-1}) \quad (2)$$

with initial condition $S(N_{0,0}) = 0$. The first term captures the expected number of new $S$-fragments generated from $M$-fragments in the preceding cycle; the second term captures the doubling of all previously accumulated $S$-fragments.

To solve the recursion, we apply it a second time by substituting $S(N_{0,c-1}) = p_2 \cdot M(N_{0,c-2}) + 2 \cdot S(N_{0,c-2})$:

$$\begin{aligned} S(N_{0,c}) &= p_2 \cdot M(N_{0,c-1}) + 2\big[p_2 \cdot M(N_{0,c-2}) + 2 \cdot S(N_{0,c-2})\big] \quad (3) \\ &= p_2 \cdot M(N_{0,c-1}) + 2\,p_2 \cdot M(N_{0,c-2}) + 4 \cdot S(N_{0,c-2}) \quad (4) \end{aligned}$$

Continuing this unrolling until the initial condition $S(N_{0,0}) = 0$ is reached gives:

$$S(N_{0,c}) = \sum_{l=1}^{c} p_2 \cdot M(N_{0,c-l}) \cdot 2^{l-1} \qquad (5)$$

Substituting $M(N_{0,c-l}) = N_0 \cdot (c-l) \cdot p_1$ (the $l = c$ term vanishes since $M(N_{0,0}) = 0$) yields:

$$S(N_{0,c}) = N_0 \cdot p_1 \cdot p_2 \sum_{l=1}^{c-1} (c-l) \cdot 2^{l-1} \qquad (6)$$

This sum can be evaluated either by applying the geometric series formula (differentiating the standard geometric sum with respect to the ratio) or via a combinatorial argument (see Appendix A.1 for both proofs). Either way, one obtains:

$$\sum_{l=1}^{c-1} (c-l) \cdot 2^{l-1} = 2^c - c - 1 \qquad (7)$$

and therefore the closed-form expression:

$$S(N_{0,c}) = N_0 \cdot p_1 \cdot p_2 \cdot (2^c - c - 1) \qquad (8)$$

For large $c$, the linear term $-(c+1)$ becomes negligible relative to $2^c$, and the expected number of short fragments grows approximately exponentially:

$$S(N_{0,c}) \approx N_0 \cdot p_1 \cdot p_2 \cdot 2^c \qquad (9)$$

This is consistent with the well-known exponential amplification behaviour of PCR.

#### 4. Amplification efficiency and relative reads

Since $p_1$ and $p_2$ act exclusively through their product in the expression for $S(N_{0,c})$, they cannot be estimated independently from sequencing data. We therefore define the combined amplification efficiency

$$E = p_1 \cdot p_2 \qquad (10)$$

which yields the final expression:

$$S(N_{0,c}) = N_0 \cdot E \cdot (2^c - c - 1) \qquad (11)$$

As the absolute number of $S$-fragments cannot be observed directly from sequencing, we work instead with relative read counts. For a sample containing $j$ distinct taxa, we define the relative reads $R_{r,i}$ of taxon $i$ as:

$$R_{r,i} = \frac{S_i}{\sum_{k=1}^{j} S_k} \quad (12)$$

Substituting $S_i = N_{0,i} \cdot E_i \cdot (2^c - c - 1)$ for each taxon, the factor $(2^c - c - 1)$ cancels, giving:

$$R_{r,i} = \frac{E_i \cdot N_{0,i}}{\sum_{k=1}^{j} E_k \cdot N_{0,k}} \quad (13)$$

This shows that the relative reads depend only on the product of amplification efficiency and initial template abundance for each taxon, and are independent of the number of PCR cycles $c$. Here, $S_i$ denotes the expected number of short fragments (Section 3); in practice, observed read counts serve as estimates of these expectations, an approximation that is highly accurate when initial copy numbers $N_{0,k}$ are large — as is typically the case in PCR-based assays (see Appendix A.3 for details).

Solving for $E_i$ yields (see Appendix A.2 for the full derivation):

$$E_i = \frac{R_{r,i} \cdot \sum_{k \neq i} E_k \cdot N_{0,k}}{N_{0,i} \cdot (1 - R_{r,i})} \quad (14)$$

Note that this expression is implicit: $E_i$ is given in terms of the efficiencies $E_k$ of all other taxa. A fully explicit solution therefore requires that the efficiencies of the remaining taxa are known — for example, by including a reference taxon with a characterised efficiency. In practice, this means $E_i$ can be estimated from observed relative reads provided that a suitable reference is available.

For all communities containing only the reference taxon and one other taxon, the efficiency of the second taxon was calculated in relation to the efficiency of the reference taxon (Table 1). These relative $E$ values were subsequently employed to predict the relative copy number from the relative reads measured after sequencing. The predicted relative copy number $R_{c,i}$ for any taxon $i$ follows directly from rearranging the expression for $R_{r,i}$. Solving for $N_{0,i}$ gives $N_{0,i} = (R_{r,i}/E_i) \cdot \sum_{k=1}^{j} E_k \cdot N_{0,k}$, where the sum is the same constant for all taxa. When forming the ratio $R_{c,i} = N_{0,i} / \sum_{k} N_{0,k}$, this constant cancels, yielding:

$$R_{c,i} = \frac{R_{r,i}/E_i}{\sum_{k=1}^{j} R_{r,k}/E_k} \quad (15)$$

This form is convenient for computation: each taxon's reads are divided by its efficiency, and the results are renormalised across all $j$ taxa. It corrects for amplification bias by down-weighting taxa with higher efficiency.

An equivalent expression, obtained by dividing numerator and denominator by $R_{r,i}/E_i$ and using $S_k/S_i = R_{r,k}/R_{r,i}$ (where $S_k$ denotes observed read counts), is:

$$R_{c,i} = \frac{1}{1 + \sum_{k \neq i}^{j} \frac{S_k\, E_i}{S_i\, E_k}} \qquad (16)$$

This form makes the pairwise structure explicit: the sum contains one term per competing taxon, each comparing the ratio of observed reads $S_k/S_i$ to the ratio of efficiencies $E_k/E_i$. If taxon $k$ amplifies more efficiently than taxon $i$ (i.e. $E_k > E_i$), its reads are proportionally discounted, increasing the estimated copy number of taxon $i$ relative to $k$. The statistical justification for using observed reads and estimated efficiencies in place of their population-level counterparts — and the consistency of the resulting estimator — is given in Appendix A.3.

#### Appendix A: Mathematical details

##### A.1 Proof of the identity used in equation (7)

We prove the identity already stated as Equation (7) in the main text, for all integers $c \geq 2$:

$$\sum_{l=1}^{c-1} (c-l) \cdot 2^{l-1} = 2^c - c - 1 \qquad (7)$$

**Proof 1: Analytic derivation**

We split the sum using linearity:

$$\sum_{l=1}^{c-1} (c-l) \cdot 2^{l-1} = c \sum_{l=1}^{c-1} 2^{l-1} - \sum_{l=1}^{c-1} l \cdot 2^{l-1}$$

**First term.** By the geometric series formula $\sum_{l=1}^{n} r^{l-1} = \frac{r^n - 1}{r - 1}$, applied with $r = 2$:

$$c \sum_{l=1}^{c-1} 2^{l-1} = c \cdot (2^{c-1} - 1)$$

**Second term.** Differentiating the geometric series $\sum_{l=0}^{n} x^l = \frac{x^{n+1}-1}{x-1}$ with respect to $x$ yields $\sum_{l=1}^{n} l \cdot x^{l-1} = \frac{d}{dx}\frac{x^{n+1}-1}{x-1}$. By the quotient rule: $\frac{d}{dx}\frac{x^{n+1}-1}{x-1} = \frac{(n+1)x^n(x-1)-(x^{n+1}-1)}{(x-1)^2}$ Evaluating at $x=2$, $n=c-1$:

$$\sum_{l=1}^{c-1} l \cdot 2^{l-1} = (c-2)\cdot 2^{c-1} + 1$$

**Combining both terms:** $c(2^{c-1}-1) - \left[(c-2)\cdot 2^{c-1}+1\right] = 2^{c-1}\left[c-(c-2)\right] - (c+1) = 2^c - c - 1$ ☐

**Proof 2: Combinatorial argument (double counting)**

**Claim.** Both sides of (A.1) count the same finite set.

Let $\mathcal{M}$ be the set of all binary strings of length $c$ that contain at least two ones. Counting directly by complement:

$$|\mathcal{M}| = 2^c - \underbrace{1}_{\text{no ones}} - \underbrace{c}_{\text{exactly one}} = 2^c - c - 1$$

We now count $|\mathcal{M}|$ again by classifying each string according to the position of its second-to-last one (i.e., the rightmost one that is not the last one). Let $l \in \{1,\ldots,c-1\}$ denote this position. For a fixed $l$, a string in $\mathcal{M}$ with second-to-last one at position $l$ is uniquely determined by:

| Segment | Constraint | Choices |
|---|---|---|
| Positions $1,\ldots,l-1$ | arbitrary bits | $2^{l-1}$ |
| Position $l$ | must be one | $1$ |
| Positions $l+1,\ldots,c$ | exactly one among $c-l$ bits | $c-l$ |

This yields $|\mathcal{M}| = \sum_{l=1}^{c-1}(c-l)\cdot 2^{l-1}$, and together with the direct count the identity (7) follows. ☐

**A.2 Derivation of the amplification efficiency $E_i$ (Equation 14)**

We derive Equation (14) from the main text in full. Starting from the expression for relative reads (Section 4):

$$R_{r,i} = \frac{E_i \cdot N_{0,i}}{\sum_{k=1}^{j} E_k \cdot N_{0,k}}$$

Multiply both sides by the denominator:

$$R_{r,i} \cdot \sum_{k=1}^{j} E_k \cdot N_{0,k} = E_i \cdot N_{0,i}$$

Separate the $k = i$ term from the sum:

$$R_{r,i} \cdot \left( E_i \cdot N_{0,i} + \sum_{k \neq i} E_k \cdot N_{0,k} \right) = E_i \cdot N_{0,i}$$

Step 3 — Expand:

$$R_{r,i} \cdot E_i \cdot N_{0,i} + R_{r,i} \cdot \sum_{k \neq i} E_k \cdot N_{0,k} = E_i \cdot N_{0,i}$$

Step 4 — Collect all $E_i$ terms on one side:

$$R_{r,i} \cdot \sum_{k \neq i} E_k \cdot N_{0,k} = E_i \cdot N_{0,i} \cdot (1 - R_{r,i})$$

Divide by $N_{0,i}(1 - R_{r,i})$, valid provided $R_{r,i} \neq 1$ and $N_{0,i} \neq 0$:

$$\boxed{E_i = \frac{R_{r,i} \cdot \sum_{k \neq i} E_k \cdot N_{0,k}}{N_{0,i} \cdot (1 - R_{r,i})}} \qquad (14)$$

Note that this result is implicit: $E_i$ is expressed in terms of the efficiencies $E_k$ of all other taxa. A fully explicit solution is only possible when the efficiencies of all remaining taxa are treated as known — for example, by designating a reference taxon with a characterised efficiency.

**A.3 Approximation in the relative read formula**

The formula derived in Section 4, Equation (13),

$$R_{r,i} = \frac{E_i \cdot N_{0,i}}{\sum_{k=1}^{j} E_k \cdot N_{0,k}} \qquad (13)$$

is obtained by substituting the expected fragment counts $\mathbb{E}[S_i]$ into the ratio. It therefore represents $\mathbb{E}[S_i]/\sum_k \mathbb{E}[S_k]$, not the expectation of the observed ratio $\mathbb{E}\left[S_i/\sum_k S_k\right]$. For random variables, these two quantities are in general not equal.

The $N_{0,i}$ starting molecules of taxon $i$ are independent and identically distributed. Denote by $X_m$ the total number of $S$-fragments produced from the starting molecule $m$ across all $c$ cycles. By the model assumptions, the $X_m$ are i.i.d. with $\mathbb{E}[X_m] = E_i \cdot (2^c - c - 1)$, and the total fragment count decomposes as:

$$S_i = \sum_{m=1}^{N_{0,i}} X_m \qquad (17)$$

The Law of Large Numbers therefore gives:

$$\frac{S_i}{N_{0,i}} \xrightarrow{P} E_i \cdot (2^c - c - 1) \quad \text{as } N_{0,i} \to \infty \quad (18)$$

Applying the same argument to every taxon $k$ independently:

$$\frac{S_i}{\sum_k S_k} = \frac{(S_i/N_{0,i}) \cdot N_{0,i}}{\sum_k (S_k/N_{0,k}) \cdot N_{0,k}} \xrightarrow{P} \frac{E_i \cdot (2^c - c - 1) \cdot N_{0,i}}{\sum_k E_k \cdot (2^c - c - 1) \cdot N_{0,k}} = \frac{E_i \cdot N_{0,i}}{\sum_k E_k \cdot N_{0,k}} \quad (19)$$

where the factor $(2^c - c - 1)$ cancels, and convergence of the ratio follows from the Continuous Mapping Theorem: the function $(x_1, \ldots, x_j) \mapsto x_i/\sum_k x_k$ is continuous wherever the denominator is non-zero.

In practice, $R_{c,i}$ is computed by plugging observed read counts $S_i/\sum_k S_k$ and experimentally determined efficiency estimates into the formula. The argument above shows that $S_i/\sum_k S_k$ converges in probability to the deterministic expression $E_i N_{0,i}/\sum_k E_k N_{0,k}$ as $N_0 \to \infty$. In the calibration experiments, all quantities except $R_{r,i}$ — namely $N_{0,i}$, $N_{0,\mathrm{ref}}$, and $E_{\mathrm{ref}}$ — are known and controlled. The efficiency $E_i$ is therefore computed as a deterministic function of the observed $R_{r,i}$ alone. Since $R_{r,i}$ is itself a random quantity (PCR is a stochastic process), so is the computed $E_i$; it is an estimator of the efficiency. By the same LLN argument as above, $R_{r,i}$ in the calibration experiment converges in probability to its deterministic counterpart as $N_0 \to \infty$, and a second application of the Continuous Mapping Theorem gives that the computed $E_i$ converges in probability to the true efficiency, and consequently that the estimate of $R_{c,i}$ converges in probability to the true relative copy number, i.e., the estimator is consistent.
